# DNA microscopy: Optics-free spatio-genetic imaging by a stand-alone chemical reaction

**DOI:** 10.1101/471219

**Authors:** Joshua A. Weinstein, Aviv Regev, Feng Zhang

## Abstract

Analyzing the spatial organization of molecules in cells and tissues is a cornerstone of biological research and clinical practice. However, despite enormous progress in profiling the molecular constituents of cells, spatially mapping these constituents remains a disjointed and machinery-intensive process, relying on either light microscopy or direct physical registration and capture. Here, we demonstrate DNA microscopy, a new imaging modality for scalable, optics-free mapping of relative biomolecule positions. In DNA microscopy of transcripts, transcript molecules are tagged *in situ* with randomized nucleotides, labeling each molecule uniquely. A second *in situ* reaction then amplifies the tagged molecules, concatenates the resulting copies, and adds new randomized nucleotides to uniquely label each concatenation event. An algorithm decodes molecular proximities from these concatenated sequences, and infers physical images of the original transcripts at cellular resolution. Because its imaging power derives entirely from diffusive molecular dynamics, DNA microscopy constitutes a chemically encoded microscopy system.

## Introduction

The spatial organization of genomes and gene products within cells and tissues is at the foundation of differentiation, specialization, and physiology in higher organisms. For example, in the central nervous system, neurons express protocadherins and neurexins in highly diverse spatial patterns that govern the cell’s intrinsic state and how it forms synapses *(1,2).* In the immune system, spatial co-localization of B- and T-lymphocytes expressing diverse immune receptors permits signaling feedback critical for immune clonal selection *(3).* In the gut, epithelial, immune, endocrine, and neural cells are spatially distributed in specific ways that impact how we sense and respond to the environment, with implications for autoimmune disease, food allergies, and cancer. In tumors, cell microenvironments may be critical for tumorigenesis *(4,5),* immune surveillance and dysfunction, invasion, and metastasis.

Although imaging of cells and tissues has been a cornerstone of biology ever since cells were discovered under the light microscope centuries ago, a gap has emerged between these methods and genomic measurements. Although both forms of measurement characterize a single biological reality, they profile the microscopic world differently. Microscopy in itself illuminates spatial detail, but does not capture genetic information unless it is performed in tandem with separate genetic assays. Conversely, genomic and transcriptomic sequencing do not inherently capture spatial details.

Recent approaches to bridge this gap rely on optical readouts that require elaborate experimental systems *(6),* physical registration and capture of molecules on grids *(7,8),* or an assumption of similarity among multiple samples so that distinct experiments performed on distinct specimens may be correlated *(9,10).* These approaches closely follow the two ways in which microscopic images have been acquired to date: (1) detecting electromagnetic radiation that has interacted with or been emitted by a sample, or (2) interrogating known locations by physical contact or ablation.

Here, we propose a novel third modality for microscopy, which requires neither optics nor physical capture from known coordinates, but relies on image reconstruction from point-proximity of individual molecules (**Fig. 1**). This principle, of determining coordinates not in relation to an absolute “Archimedean point” but instead in relation to one another, features prominently in the theory of sensor localization *(11)*. Numerical work has further shown that sparse and noisy measurements of pairwise distances between points can be used to reconstruct their positions *(12).* We build on this theoretical concept to demonstrate a novel form of microscopy, called DNA microscopy. DNA microscopy reformulates the point-localization problem by reconstructing the positions of molecules using the stochastic output of a stand-alone chemical reaction. We confirm that DNA microscopy is able to resolve the physical dimensionality of a specimen, and then demonstrate that it is able to accurately reconstruct a multicellular ensemble *de novo* without optics or any prior knowledge on the organization of biological specimens.

**Fig. 1.**
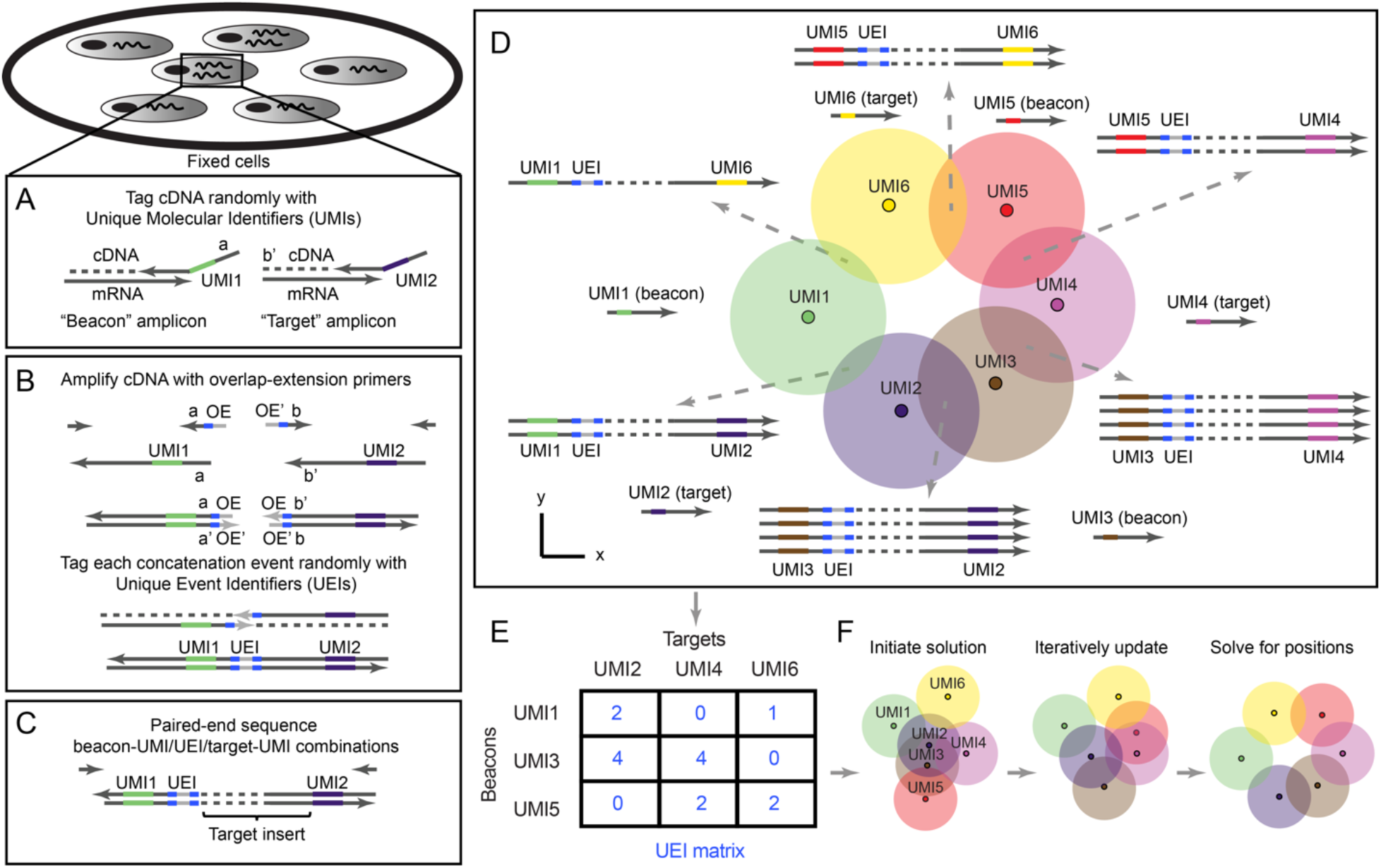
DNA microscopy. **(A-B)** Method steps. Cells are fixed and cDNA is synthesized for beacon and target transcripts with randomized nucleotides (UMIs), labeling each molecule uniquely **(A)**. *In situ* amplification of UMI-tagged cDNA directs the formation of concatemer products between beacon and target copies **(B)**. The overhang-primers responsible for concatenation further label each concatenation event uniquely with randomized nucleotides, generating unique event identifier (UEIs). Paired-end sequencing generates read-outs including a beacon-UMI, a target-UMI, the UEI that associates them, and the target gene insert **(C)**. A bird’s-eye view of the experiment **(D)** shows the manner in which the DNA microscopy reaction encodes spatial location. Diffusing and amplifying clouds of UMI-tagged DNA overlap to extents that are determined by the proximity of their centers. UEIs between pairs of UMIs occur at frequencies determined by the degree of diffusion cloud overlap. These frequencies are read out by DNA sequencing, and inserted into a UEI matrix **(E)** that is then used to infer original UMI positions **(F)**.

Finally, we demonstrate its ability to resolve and segment individual cells for transcriptional analysis.

## Results

### DNA microscopy for spatio-genetic imaging

Intuitively, DNA microscopy generates images by first randomly tagging individual DNA or RNA molecules with DNA-molecular identifiers. Each deposited DNA-molecular identifier then “communicates” with its neighbors through two parallel processes. The first process broadcasts amplifying copies of DNA-molecular identifiers to neighbors in its vicinity via diffusion. The second process encodes the proximity between the centers of overlapping molecular diffusion clouds: DNA-molecular identifiers undergo concatenation if they belong to diffusion clouds that overlap. Finally, an algorithm infers from these association rates the relative positions of all original molecules.

DNA microscopy is premised on the notion that DNA can function as an imaging medium in a manner equivalent to light. In the same way that light microscopy images molecules that interact with photons (either due to diffraction or scattering or because these molecules emit photons themselves) and encodes these images in the wavelengths and directions of these photons, DNA microscopy images molecules that interact with DNA (including DNA, RNA, or molecules that have been tagged with either DNA or RNA) and encodes these images in the DNA sequence products of a chemical reaction.

With this analogy in mind, we can imagine superposing two distinct physical processes: a fluorophore radially emitting photons at a specific fluorescence wavelength, and a DNA molecule with a specific sequence undergoing PCR amplification, and its copies diffusing radially. Optical microscopes use lenses to ensure that photons hitting a detector or the human eye will retain some information, based on where they hit, regarding their point of origin. However, the “soup” of DNA molecules generated in a DNA microscopy reaction does not afford this luxury. We therefore need a different way to distinguish the identities of point sources so that all data is encoded into the DNA itself.

To molecularly distinguish point sources we rely on Unique Molecular Identifiers, or UMIs *(13),* consisting of randomized bases that tag a molecule before any copy of it has been made (**Fig. 1A**). Because the diversity of UMIs scales exponentially with their length, we have high confidence that when one long UMI tags a molecule, no other molecule in the rest of that sample has been tagged with that same long UMI. We can now use overlap extension PCR to concatenate the diffusing and amplifying copies of these UMIs (with any biological DNA sequences they tag simply carried along). The rate at which they concatenate will reflect the distance between their points of origin.

However, once we sequence the final DNA products, we are still left with the problem of how to quantitatively read out these concatenation rates from DNA sequence alone. Using read-abundances belonging to concatenated DNA products carries serious drawbacks. For example, trace cross-contamination between samples could easily introduce artifactual UMI-UMI associations, and biases in downstream DNA library preparation could heavily distort association frequencies. Most serious, however, would be PCR chimerization: any *ex situ* amplification of the DNA library would necessarily introduce template-switching at some rate that would corrupt the data.

We reasoned that if the overlap extension primers contained randomized bases that did not participate in priming themselves, then although each priming event would result in replacement of this randomized sequence, each overlap extension event would fix the new bases in between the now-concatenated sequences (**Fig. 1B**). The concatenated sequences would then carry these randomized bases forward, intact, as they amplified. These bases would from then on be a unique record of that *individual concatenation event.* We called these new concatenated randomized sequences Unique Event Identifiers, or UEIs, and used them to encode molecular positions into the DNA microscopy reaction.

### Experimental assay for DNA microscopy to encode relative positions of molecules in cells

To demonstrate DNA microscopy, we aimed to image transcripts belonging to a mixed population of two co-cultured human cell lines, GFP-expressing MDA-MB-231 cells and RFP-expressing BT-549 cells. We reasoned that an initial proof of concept would be to recover images that appear cell-like and where GFP and RFP transcripts are positioned in mutually-exclusive cells, whereas GAPDH and ACTB, expressed in both cell lines, are ubiquitous.

In the first step of the experiment, we tag with Unique Molecular Identifiers (UMIs) cDNA synthesized *in situ*. We designed reaction chambers to both grow cells and perform all reactions (**Fig. S1, Supp. Info**.). We cultured the cells, and, following fixation and permeabilization, synthesized cDNA by reverse transcription from GFP, RFP, GAPDH, and ACTB gene transcripts (**Tables S1–S2**), with primers tagged with 29nt long UMIs (**Fig. 1A, Fig. S2**). Notably, we designed the reaction to distinguish two types of UMI-tagged cDNA molecules: “beacons”, synthesized from ACTB (chosen as a universally expressed gene whose sequence would not be analyzed in later stages), and “targets” (everything else). We achieved this distinction between beacon and target amplicons by the artificial sequence-adapters assigned to the primers annealing to each.

In the second step of the experiment, we allow beacon-cDNA and target-cDNA molecules, along with the UMIs that tag them, to amplify, diffuse, and concatenate *in situ* in a manner that generates a new Unique Event Identifier, or UEI, distinct for each concatenation *event* (**Fig. 1B and S2**) through overlap-extension PCR *(14).* By design, target amplicon-products will only concatenate to beacon amplicon-products, thereby preventing self-reaction, and the middle of each overlap-extension primer includes 10 randomized nucleotides, such that each new concatenation event generates a new 20-nucleotide UEI. Paired-end sequencing of the final concatenated products generates reads each containing a beacon UMI, a target UMI, and a UEI associating them (**Fig. 1C**).

The key to DNA microscopy is that because UEI formation is a second order reaction involving two UMI-tagged PCR amplicons, *UEI counts* are driven by the co-localization of UMI concentrations, and thus contain information on the proximity between the physical points at which each UMI began to amplify (**Fig. 1D**). In particular, as UMI-tagged cDNA amplifies and diffuses in the form of clouds of clonal sequences that overlap to varying extents, the degree of overlap (**Fig. 1D**, circle intersection) – and thus the probability of concatenation and UEI formation – depends on the proximity of the original (un-amplified) cDNA molecules (**Fig. 1D**, small dark circles). UMI-diffusion clouds with greater overlap generate more UEIs/concatemers, whereas those clouds with less overlap generate fewer UEIs/concatemers.

To obtain reliable estimates of UEIs between every pair of UMIs, we must address sources of noise, such as sequencing error. We cluster beacon-UMIs, target-UMIs, and UEIs by separately identifying “peaks” in read-abundances using a log-linear time clustering algorithm (**Supp. Info., Fig. S3A**) in a manner analogous to watershed image segmentation, but in the space of sequences. For target UMIs, this allows us to aggregate biological gene sequences originating from *single* target molecules and achieve low error rates by taking a consensus of the associated reads (**Fig. S3B**). We then assign each identified UEI a single consensus beacon-UMI/target-UMI pair based on read-number plurality, and prune the data (by eliminating UMIs associating with only one UEI) to form a sparse matrix whose elements contained integer counts of UEIs pairing each beacon-UMI (matrix rows) and each target-UMI (matrix columns) (**Fig. 1E, Supp. Info**.). The resulting UEI matrices, containing on the order 10^5^ -10^6^ total UMIs, and averaging ~10 UEIs per UMI (**Table S3**), form the data sets upon which we built an engine for image inference.

### A “zoom” function infers local spatial encodings from UEI matrices

Next, we developed an algorithmic approach to use UEI prevalence to infer UMI proximity and reconstruct an image of the original sample and its transcripts (**Fig. 1F**). We first appreciate that if the UEI matrix had successfully encoded relative UMI coordinates, these coordinates would be reflected in the rows and columns of the matrix. The matrix rows and columns would span a space having a dimensionality scaling with the total number of UMIs. However, if they encoded UMI coordinates within a sample, they would collectively sweep out a far smaller dimensionality, only equal to the physical dimensionality of the sample.

As a toy example, consider a comparison between three systems in which a single target UMI (“2”) is in each of three positions in one dimension relative to two beacon UMIs (“1” and “3”) with which it forms UEIs (**Fig. 2A**). The target UMI begins closest to one of the two beacon UMIs, and as a result, its diffusion cloud overlaps most with that beacon UMI’s diffusion cloud. Thus, its reaction rate with that beacon UMI is relatively higher (**Fig. 2B**) and results in a correspondingly larger number of UEIs (**Fig. 2C**). If the target UMI is further away, the balance of overlaps between diffusion clouds changes. Indeed, plotting expected UEI matrix elements for the target UMI on two axes, we see that its trajectory remains onedimensional (**Fig. 2D**). Extending to a large population of target UMIs across many positions, these new target UMIs, just like the target UMI in the original example, also interact with the same two beacon UMIs. Therefore, we can also plot them on the same two axes, and wherever they land, we could expect them to scatter around the same onedimensional manifold followed by the target UMI of the original example. In any real data set, UEI count is affected not only by position but also by additional variables (such as amplification biases and diffusion rates), each potentially adding to the data’s total dimensionality. However, these sources of variation would be *suppressed along the principle dimensions* of a UEI matrix so long as their effect on neighboring UMIs is not systematically correlated.

**Fig. 2.**
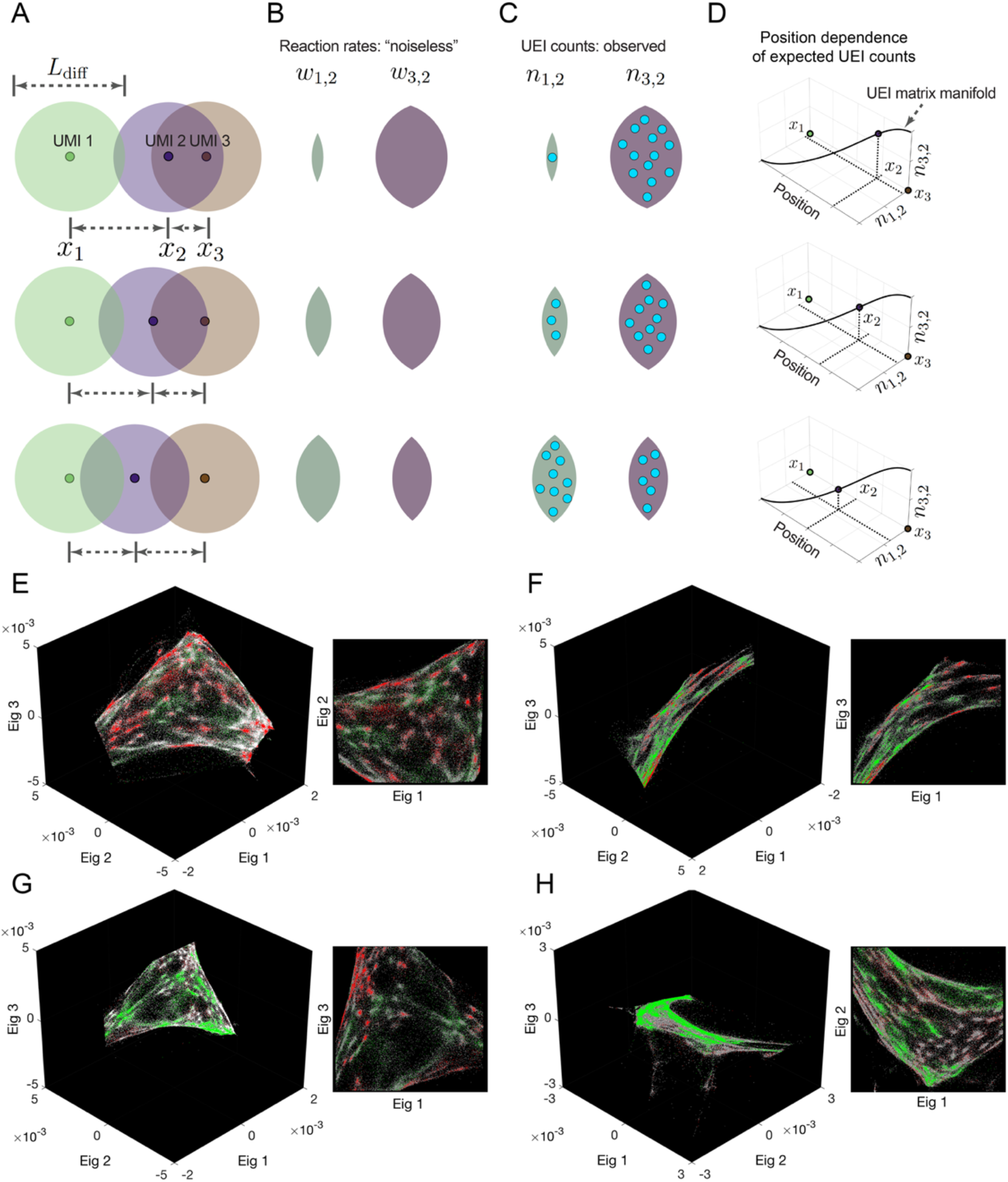
Encoding and decoding molecular localization with DNA microscopy. Diffusion profiles with length scale *Ldiff* belonging to different amplifying UMIs overlap to degrees that depend on the distance between their points of origin **(A)**. Greater overlaps between diffusion profiles result in larger reaction rates **(B)**, which in turn result in higher UEI formation frequencies **(C)**. Because UEI counts are therefore proper functions of position, as a UMI relocates it sweeps out a curve along the UEI count axes equal to the dimensionality of space it occupies **(D)**. **(E-H)** Data segmentation permits individual sets of 10^4^ strongly interacting UMIs to be visualized independently. The top three non-trivial eigenvectors for the largest data segments of samples 1 and 2 are shown, along with a different, magnified view of the same plot. Transcripts are colored by sequence identity: grey = ACTB (beacons), white = GAPDH, green = GFP, red = RFP.

To identify the principle dimensions of the UEI matrix, we can analyze the graph of UMI vertices and weighted UEI count edges by constructing a Graph Laplacian matrix from the raw UEI matrix (with its diagonal elements set so that each row sums to zero). The Graph Laplacian eigenvectors with the smallest-magnitude eigenvalues would visualize the most systematic forms of variation in the DNA microscopy data (**Supp. Info**.) and illuminate the low-dimensional manifold, if any, it occupied. However, even a low-dimensional manifold could be folded in complex ways in the high-dimensional space formed by a full UEI matrix, making it difficult to analyze the manifold’s shape over large distances, especially in areas of the manifold that were sparsely populated. Analyzing the UEI matrix manifold therefore first requires analyzing UMI subsets corresponding to *local* regions of the original sample. We return to global relations in subsequent sections.

To perform this local investigation, we developed a “zoom” function for DNA microscopy data by applying a recursive graph-cut algorithm, identifying putative cuts by using the spectral approximation to the cut of minimum-conductance *(15)* (**Supp. Info**.). This criterion separates sub-sets of UMIs exhibiting UEI-flux between them that was relatively small given the number of UMIs they comprised. The algorithm first finds the sparsest cut to the entire data set, then the sparsest cuts to the resulting halves, and so on until a further sparse cut cannot be made (**Supp. Info.**). We then visualize each of these sub-regions by the eigenvectors corresponding to the smallest-magnitude eigenvalues of their UEI-Graph Laplacian sub-matrix.

### Successful inference of local structure identifies cell-like structures with specific marker expression

Strikingly, and consistent with our theoretical reasoning, although the UMIs in these subsets fully spanned at least all three eigenvector dimensions, the manifolds swept out by the UMIs were only two-dimensional (Fig. 2E-H). This confirmed that neighborhoods of points had been successfully encoded into the UEI matrix: the dimensionality of their spatial relationships within the sample was correctly preserved.

The two-dimensional manifolds exhibited clusters of UMIs that recapitulated the genetic composition of the cell lines used in the experiment: a pervasive distribution of the constitutively-expressed ACTB and GAPDH sequences, but a mutually exclusive distribution of GFP and RFP, recapitulating their correct cell specific expression (**Fig. 2E-H**). An observer unaware of the spatial dimensionality of the specimen or that cells even existed could discover both by analyzing the DNA microscopy sequencing data alone. Together, these two observations confirmed both cellular and local supra-cellular resolution in DNA microscopy.

### Inference of global molecular positions from DNA microscopy data

Next, we expanded our inference beyond the local scope of a few thousands of proximal transcript molecules, by developing a framework for evaluating the likelihood of a global position-estimate solution.

We reasoned that each UEI’s occurrence is analogous to a “coin-toss” experiment performed on every UMI-pair, with each pair’s “meeting” probability proportional to the corresponding reaction rate (**Fig. 3A, Supp. Info**.). In this probability function, UEIs in DNA microscopy act in the same manner as photons do in optical super-resolution localization microscopy *(16):* both narrow a point-spread function governed by a physical length scale (wavelength in the case of light, diffusion distance in the case of DNA) as they accrue in number (**Fig. 3B,C**). In real data sets, UEIs increase progressively with increasing read depth, whereas UMIs saturate more quickly (**Fig. 3D,E**). In this way, read depth in DNA microscopy constitutes a dial to increase the number of UEIs per UMI, enhancing an image’s resolution.

**Fig. 3.**
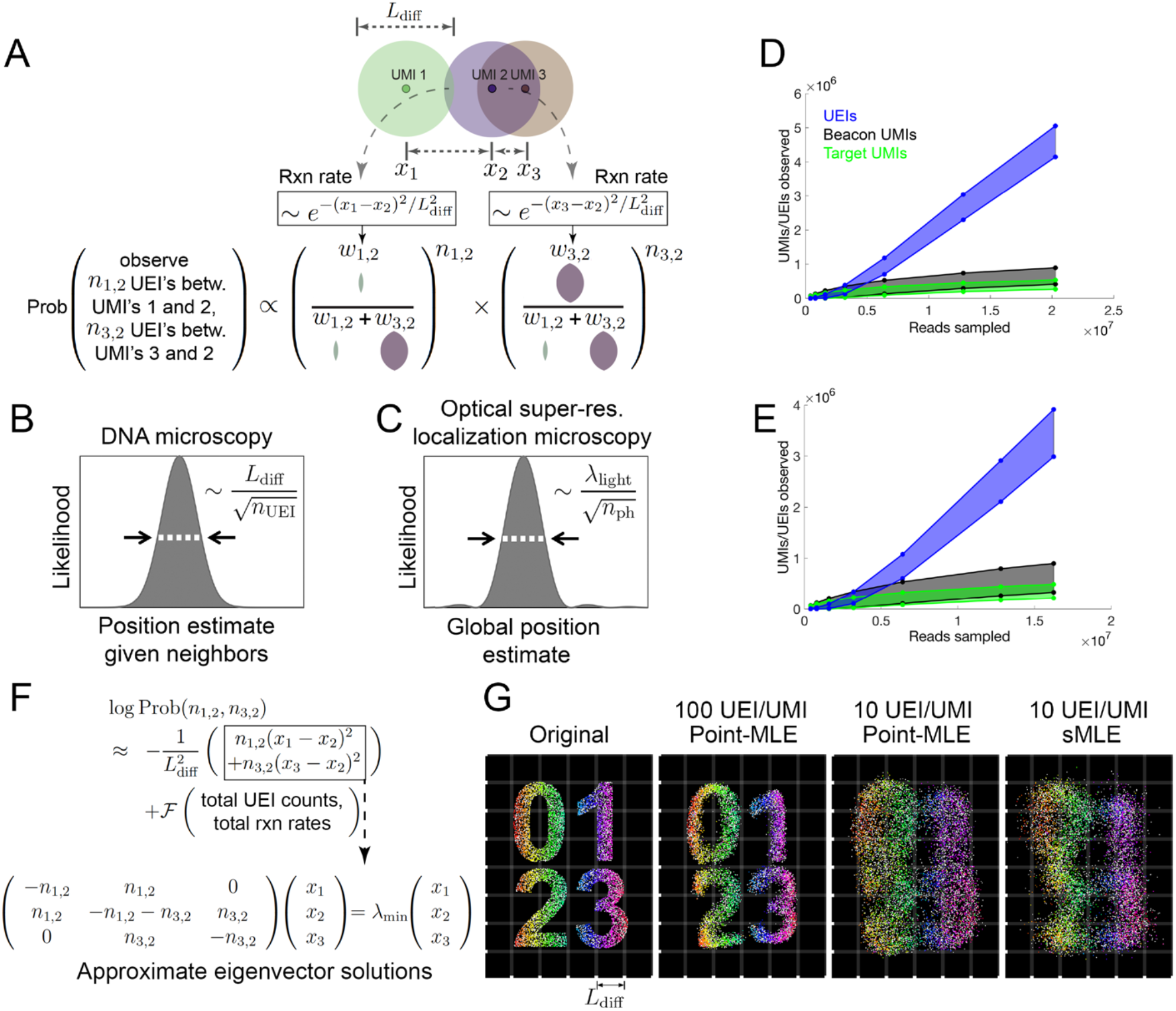
Image inference from DNA microscopy data. **(A)** Modeling diffusion of amplifying UMIs as isotropic across length scale *L_diff_* allows the likelihood of a UMI-position solution to be evaluated given observed UEI counts. **(B,C)** Uncertainty in DNA vs. super-resolution microscopy. Given its reacting partners’ positions, DNA microscopy (left) defines a UMI’s uncertainty as a physical length-scale (DNA diffusion distance, *L_diff_*) divided by the square-root of the number of individual quanta measured (UEIs) in a manner analogous to quanta (photons) in super-resolution microscopy (right). **(D,E)** Rarefaction of UMI and UEI data. Shown are curves with an upper-bound, indicating total UMI/UEI counts, and a lower-bound, indicating those from the final pruned UEI matrix, for samples 1 **(D)** and 2 **(E)**. **(F)** The sMLE algorithm uses eigenvector solutions to part of the position-probability function to identify a linear basis for the solution to the full likelihood function. **(G)** sMLE enhances performance in free-diffusion simulation tests. From left: original image, results from point-MLE on simulated images with 100 or 10 UEIs/UMI, and from sMLE with 10 UEIs/UMI.

Unlike its optical counterpart, however, DNA microscopy resolves a molecule’s position by orienting it relative to other molecules, and its uncertainty is therefore a function of these relationships. A relationship between two UMIs may come in two forms: those that are *direct* and involve UEIs linking them, and *indirect* relationships that occur via intermediaries. The latter emerges in the structure of the data, but will not strongly influence UMI positions if these positions are optimized independently. This may be seen in the logarithm of the UEI-count probability function (**Fig. 3F**). This log-probability is the sum of two components: (1) a sum of squared-differences between positions, weighted by individual UEI counts; and (2) a function of total UEI counts and total expected reaction rates (which are themselves functions of UMI positions) across the entire data set. In order to still calculate the log-probability as a whole in a scalable way, we implemented the Fast Gauss Transform *(17)* (**Fig. S6**).

If each UMI’s position is updated independently to maximize this log-probability function, it will experience two forces, corresponding to the function’s two added components: the first which pulls together UMIs that have directly formed UEIs between them, and the second which repels all UMIs from all other UMIs. The likelihood of the position-solution is maximized when these two forces balance. During the maximization update-process, the only way in which an *indirect* relationship between UMIs will influence their position-solution is if intermediary UMIs that *directly* form UEIs with them separately have already changed position.

To ensure that large-scale optimization captures these indirect UMI relationships encoded in the data, we developed a new maximum likelihood framework, which we called spectral maximum likelihood estimation or sMLE, to generate global representations of the DNA microscopy data. Because maximizing the first component of the log-probability entails minimizing the magnitude of the sum of squared-differences, it can be individually solved by identifying the smallest-magnitude eigenvalue/eigenvector pairs of the UEI Graph Laplacian introduced earlier (**Fig. 3F, Supp. Info**.). Each eigenvector represents a distinct way in which UMIs can be *globally* rearranged to suit orientation requirements expressed by the sum of squared-differences between *local* points. The eigenvector with the smallest-magnitude eigenvalue represents the best arrangement, the second smallest-magnitude eigenvalue the second best, and so on. Because sums of eigenvector solutions to the local linear problem would produce solutions that themselves satisfy local constraints, sum-coefficients of these eigenvectors could act as variables in a larger-scale likelihood maximization. By seeding a solution with the two eigenvectors corresponding to the smallest-magnitude eigenvalues, optimizing their coefficients, then incorporating successive eigenvectors and repeating, we could find global solutions that were also well-constrained locally. These sMLE solutions showed strong advantages in simple simulations over maximizing the likelihood while treating every UMI independently, especially when UEI counts were limiting (**Fig. 3G**).

### DNA microscopy correctly recapitulates optical microscopy data

We next sought to apply the sMLE inference framework to determine whether DNA microscopy could resolve supra-cellular coordinates compared to optical microscopy. To this end, we constructed reaction chambers with glass slides (**Fig. S1B**) and plated GFP-and RFP-expressing cells in a highly localized pattern within the chamber (**Fig. S1C**). We then imaged GFP- and RFP-expression in cells across the entire area of the reaction chamber using an epifluorescence microscope before the DNA microscopy reaction (**Fig. 4A,B**), sequenced the resulting DNA library to saturation (**Fig. 4C**), and applied the sMLE inference algorithm.

**Fig. 4.**
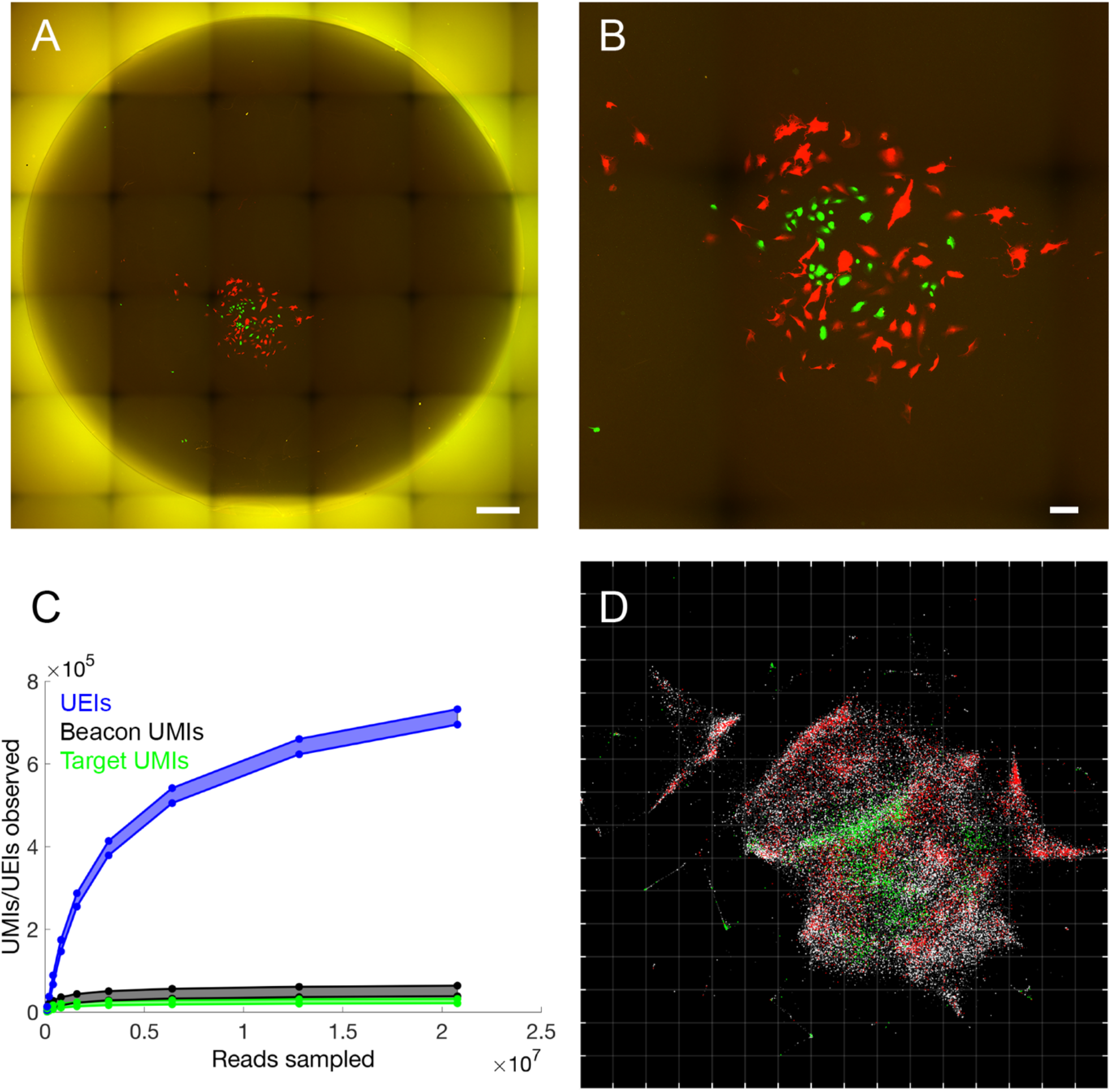
Accurate reconstruction by DNA microscopy of fluorescence microscopy data. **(A)** Full reaction chamber view of co-cultured GFP- and RFP-expressing cells (scale bar = 500 um). **(B)** Zoomed view of the same cell population (scale bar = 100 um). **(C)** Rarefaction of UMIs and UEIs with increasing read-sampling depth. **(D)** sMLE inference applied to DNA microscopy data, reflected/rotated and rescaled for visual comparison with photograph. Transcripts, sequenced to 98 bp, are colored by sequence identity: grey = ACTB (beacons), white = GAPDH, green = GFP, red = RFP. Grid-line spacings: diffusion length-scales (*L_diff_*), emerging directly from the optimization (**Supp. Info**.).

Strikingly, the resulting image recapitulates optical microscopy data without systematic distortion (**Fig. 4D**) in both the shape of the cell population boundary, as well as the distribution of GFP- and RFP-expressing cells within it. Importantly, the inferred image preserves the correct aspect ratio: although needing to be rotated and reflected, the individual axes did not need to be independently re-scaled. This demonstrated that DNA microscopy is capable of generating accurate physical images of cell populations.

### Large length-scale optimization and the folded manifold problem

We next sought to apply DNA microscopy to optimization at larger length scales. Applying sMLE inference to the original data generated from several hundred cells used to generate the original eigenvector representations (**Fig. 2**) gave images that reproduced the individual cell compositions of the earlier visualizations (**Fig. 5**). These large-scale optimizations were also robust to data down-sampling (**Fig. S4**). Nevertheless, the reconstructed images exhibited “folding” that indicated how the process of projecting large and curved high-dimensional manifolds onto two-dimensional planes was vulnerable to distortions. The causes for this “manifold folding” problem could be understood by examining the illustration of how low-dimensional manifolds come into being within a high-dimensional UEI data matrix (**Fig. 2**). Eigenvector calculation (**Fig. 3F**) involves identifying hyperplanes that can be drawn through these low-dimensional manifolds that maximally account for variation in the UEI data. It does this in a manner similar to linear regression, balancing the advantage of fitting certain parts of the data with the costs of not fitting other parts of the data.

**Fig. 5.**
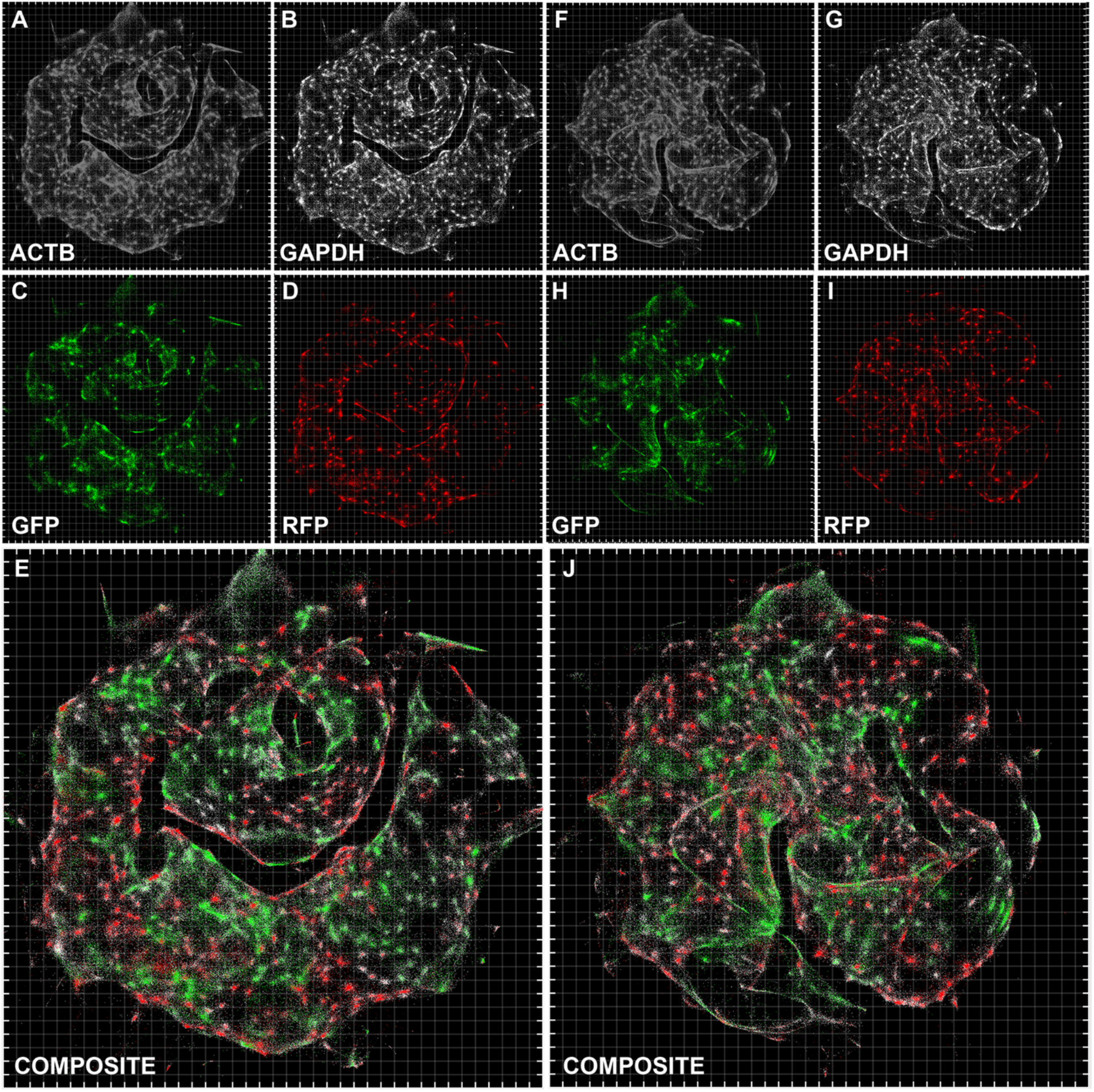
Inferred large scale DNA microscopy images preserve cellular resolution. Inference using the sMLE global inference approach for sample 1 **(A-E)** and sample 2 **(F-J)**, with each transcript type shown separately **(A-D, F-I)** or together **(E and J)** (although inferences are performed on all transcripts simultaneously and are blinded to transcript identity). Grid-line spacings: diffusion length-scales (*L_diff_*) emerging directly from the optimization (**Supp. Info**.).

However, this balancing can yield errors in several ways. If a large number of UMIs in one part of the data set rotate the top calculated eigenvectors (with the smallest-magnitude eigenvalues) away from UMIs in a different part of the data set, then projecting the global data set onto these eigenvectors will cause these neglected UMIs to fold on top of one another. This will produce the type of artifact observed for large scale optimization (**Fig. 5**). If we avoid eigenvector calculation entirely and optimize each UMI’s position independently (**Fig. S5A,B**) we avoid such defects, but obtain close-packed images, as predicted by simulation (**Fig. 3G**), that do not preserve empty space. This highlights the distinct nature of DNA microscopy’s imaging capabilities compared to light’s, where density rather than sparsity is the key challenge.

### Cell segmentation can be performed on the UEI matrix based on diffusion distance

We next analyzed the degree to which the UEI matrix could be used to segment cells and analyze single cell gene expression. Importantly, up to this point, no step in the process – experimental or computational – had knowledge that cells even exist. To perform segmentation, we applied the same recursive graph cut algorithm as used earlier to generate local eigenvector visualizations of the data. By increasing the conductance-threshold dictating whether segments of the data should be left intact, we assigned transcripts to putative cells (Fig. 6A,B), again without regard to transcript identity (i.e., GFP vs. RFP). To quantify segmentation quality, we calculated the probability that, within each putative cell, the minority fluorescent gene transcript would occur at or lower than its current value, given its prevalence in the data set. We found the median p-value decayed rapidly, over a range of conductance thresholds, to <10^−10^, with increasing reads and resolved cells analyzed (**Fig. 6C,D**).

**Fig. 6.**
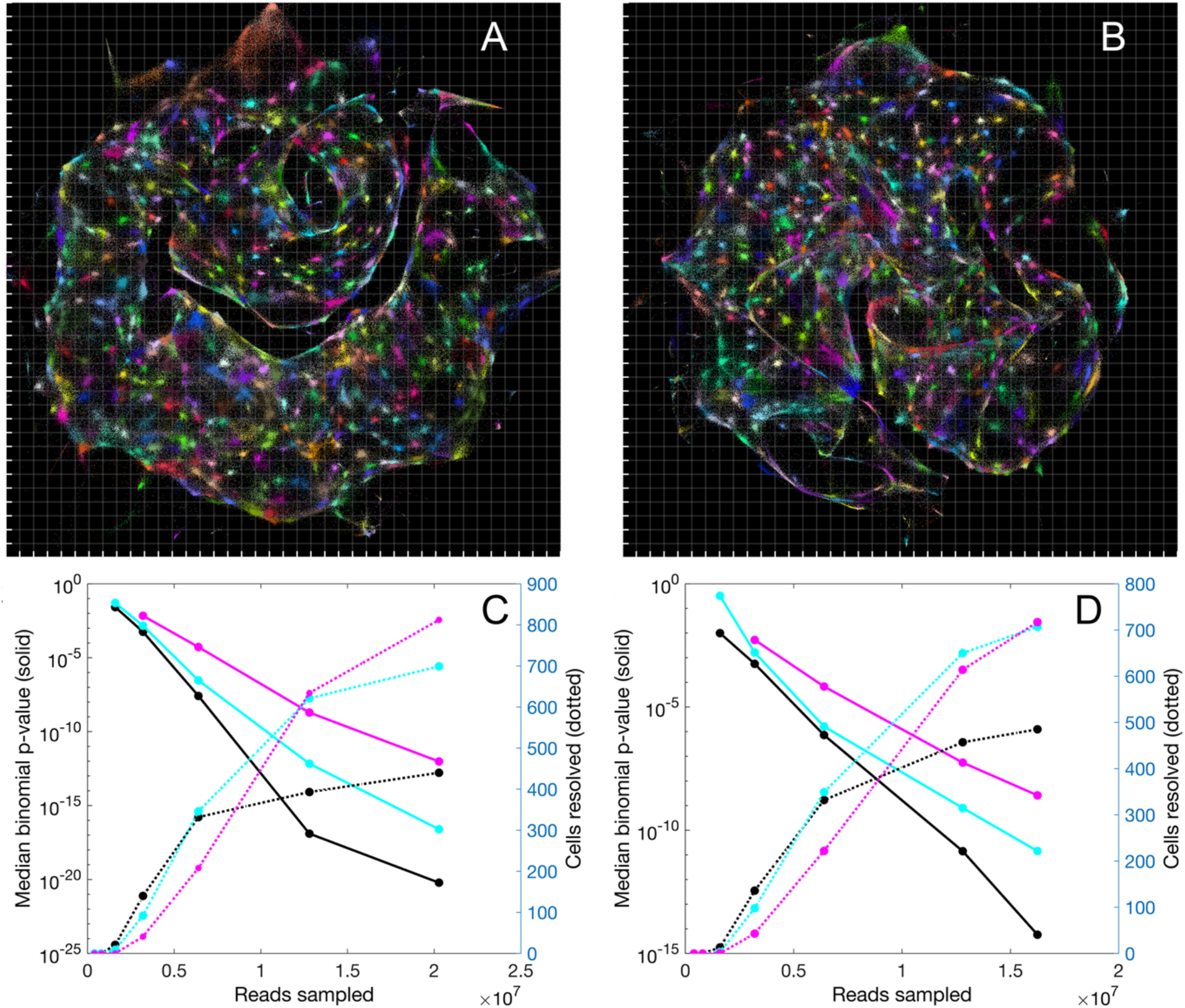
Segmentation of DNA microscopy data recovers cells *de novo.* **(A,B)** Data segmentation recovers putative cells without a priori knowledge. Cell segmentation for samples 1 **(A)** and 2 **(B)** by recursive graph-cutting of the UEI matrix is shown with a random color assigned to each inferred cell, qualifying if it contained at least 50 UMIs and had at least one transcript each of ACTB and GAPDH. The minimum conductance threshold was set to 0.2. **(C,D)** Segmentation performance. The effects of cell segmentation for samples 1 **(C)** and 2 **(D)** with minimum conductance thresholds 0.14 (black), 0.2 (cyan), and 0.26 (magenta) are shown on binomial p-values quantifying segmentation fidelity (solid lines) and putative cell count (dotted lines).

### Imaging large numbers of different transcripts in DNA microscopy

To demonstrate that DNA microscopy and its associated cell segmentation could be extended to larger numbers of genes, we synthesized cDNA by reverse transcription from up to 20 additional genes that were known to be differently enriched in MDA-MB-231 and BT-549 cell lines (**Tables S4–S5, Supp. Info**.). We performed global image inference (**Fig. S5**) and applied the same hierarchical cell segmentation algorithm as earlier. Pearson correlations between GFP-fraction (out of total transgene transcripts per cell) and fraction of endogenous genes expected enriched in the GFP cell line (out of total endogenous gene transcripts enriched in either cell line) gave r = 0.29-0.41 (n=764 and 265) for two experiments, respectively (p-value<10^−6^, permutation test). This demonstrated that the transgenes labeling these cell types retained information about cell type-specific endogenous expression, and that this information could be read out from DNA microscopy data.

## Discussion

The fundamental advance of DNA microscopy is to physically image biological specimens using an unstructured and standalone chemical reaction. This makes it a distinct microscopic imaging modality in itself. As a technology, we have drawn a close parallel between DNA microscopy and optical super-resolution: both take advantage of stochastic physics to reduce measurement uncertainty beyond what may seem superficially to be a limit imposed by physics.

However, the two differ in several fundamental ways. Optical super-resolution microscopy relies on the quantum mechanics of fluorescent energy decay. DNA microscopy, however, relies entirely on thermodynamic entropy. The moment we tag biomolecules with UMIs in the DNA microscopy protocol, the sample gains spatially-stratified and chemically-distinguishable DNA point sources. This process thereby introduces a spatial chemical gradient across the sample that did not exist previously. Once these point sources begin to amplify by PCR and diffuse, this spatial gradient begins to disappear. This entropic homogenization of the sample is what drives different UMI diffusion clouds to interact and UEIs to form. It is therefore this increase in the system’s entropy that most directly drives the DNA microscopy reaction to record meaningful information about a specimen.

One key point of weakness for DNA microscopy remains the resolution of empty space, and future work will be needed to eliminate this obstacle to produce high quality reconstructions of samples over large lengths where there are gaps. It is possible that a “landmark” based approach, in which specific DNA sequences are deposited at known physical locations to assist in the image reconstruction process, will ultimately prove the most cost-effective way to achieve this. Better analytical techniques to correct for large length scale distortions may prove equally effective, without complicating the experiment itself. Nevertheless, the fact that DNA microscopy is performed entirely by pipette means that large numbers of samples can easily be processed simultaneously. The technology is therefore structurally conducive to massive throughput.

DNA microscopy offers a new form of optics-free imaging that leverages the large economies of scale in DNA sequencing. The technology does not require sacrificing spatial resolution for sequence accuracy, since it benefits, rather than suffers, from high signal density and it does not hinge on optical resolution of diffraction-limited “spots” *in situ*. By using chemistry itself as its means of image acquisition, DNA microscopy decouples spatial resolution from specimen penetration depth (otherwise linked by the properties of electromagnetic radiation) and thereby side-steps a tradeoff imposed by the physics of wave propagation.

Finally, because it does not rely on specialized equipment and can be performed in a multi-well format with normal lab pipettes, DNA microscopy is highly scalable. It is fully multiplex-compatible (imaging any PCR template) and uses sequencing-depth as a dial to enhance spatial and genetic detail. Moreover, because DNA microscopy reads out single-nucleotide variation in biological DNA or RNA sequences it targets, it spatially resolves the astronomically large potential variation that exists in somatic mutations, stochastic RNA splicing, RNA editing, and similar forms of genetic diversity in cell populations. We have demonstrated that it achieves this at high accuracy over long read lengths.

Our development of a chemically-encoded microscopy system leaves open both fundamental theoretical and experimental questions. On the one hand, future experimental and computational enhancements will help better resolve large length scales that include spatial gaps. On the other hand, the UEI, by effectively functioning in these experiments as a DNA-analogue of the photon, has illuminated a wider potential role for DNA as a medium for artificial biological recordings. Most directly, DNA microscopy can be applied in principle beyond the transcriptome, for example, directly to DNA sequences or to proteins detected with DNA-labeled antibodies. Looking to the future, a full exploration of individual and idiosyncratic spatial structures in the biological world by encoding them into DNA bases, instead of photonic pixels, may reveal new layers of information otherwise hidden by the limits of optical- and electron-based imaging.

## Acknowledgments

We thank Paul Blainey and his lab for their generosity and provision of their clean room for PDMS work, as well as for particularly helpful discussions with David Feldman and Lily Xu.

## Author contributions

J.W. conceived the project. J.W. performed all experiments and analysis. J.W., A.R., and F.Z. wrote the manuscript.

## Supplementary Information

### 1 Experimental methods

#### 1.1 Bead-plate reaction chambers

Reaction chambers for large cell populations in samples 1-2 and 4-5 (**Fig. S1A**) were designed in order to maximally adhere cells while providing a thermally robust container for PCR thermocycling. 3 mm glass beads (Sigma Z265926) were acid washed in 1 M HCl at 50-60 C for 4-5 hours in a glass beaker with occasional agitation and then kept sealed in 90% ethanol at room temperature until further use. After an initial rinse in acetone, beads were treated for 60 seconds in a 2% solution of (3-aminopropyl)triethoxysilane/APTES (Sigma 440140) in acetone. Beads were then rinsed 4 times in ddH_2_O, rinsed in isopropanol, and allowed to dry in a laminar flow hood 1 hour in a polystyrene petri dish. Dried beads were kept sealed at room temperature until further use.

PDMS (R.S. Hughes RTV615) was mixed at a ratio of 1:10 w/w cross-linker:potting reagent, and mixed/degassed 3 minutes at 2000 rpm. Uncured PDMS was immediately dispensed into PCR plate wells (Axygen PCR-96-HS-C) at ~ 20 ul in volume. Plates were then spun down at 500×g for 1 minute. Volumes were carefully equalized across wells, and the plate was spun down again. APTES-treated glass beads were then placed into each PDMS-filled well of the PCR plate using plastic tweezers. The PCR plate was then spun down again at 500×g for 5 minutes, and beads were checked to ensure a small amount of surface was exposed above the PDMS. Bead-plates (illustrated in **Fig. S1A**) were then cured at 80 C for 2 hours, and stored sealed at room temperature until further use.

#### 1.2 Glass-slide reaction chambers

Reaction chambers for imaged cells in sample 3 are shown in **Fig. S1B-C**. PDMS was mixed at a ratio of 1:10 w/w cross-linker:potting reagent as before, and mixed/degassed 3 minutes at 2000 rpm. Uncured PDMS of mass 33-35 g was immediately dispensed into 10 cm petri dishes and degassed under vacuum for 1 hour. PDMS was then cured at 80 C for 150 minutes, and holes were punched using Integra biopsy punches with diameter 6 mm in the pattern indicated (**Fig. S1B**). Cut PDMS blocks were then bonded with oxygen plasma to plain glass slides (VWR 16004-422) and cured at 80 C for 3 hours. 100-120 ul of mineral oil (Sigma M5904) was then added to all wells and degassed 45 minutes. Slides were then baked at 80 C for 5 hours, then allowed to cool, and mineral oil was aspirated. Slides were washed heavily with acetone and isopropanol to get rid of residual mineral oil and allowed to dry. 2% APTES solution was prepared in acetone as above, and the bottom of the wells were immersed with 35 ul of this solution for 60 seconds. Wells were immediately rinsed 5 times with 120 ul water, 2 times with isopropanol, allowed to dry, and stored sealed at room temperature until further use.

#### 1.3 Cell seeding

Before cell seeding, bead-plates were rinsed twice with 70% EtOH and allowed to dry 45 minutes under UV in a cell culture hood. All wells were then washed once with 100 ul DPBS (Sigma D8537). A fibronectin solution (Sigma F1141) was then prepared at a 1:100 dilution in DPBS and used to cover wells, which were left at room temperature for 1 hour. BT-549-RFP (Cell Biolabs AKR-255) and MDA-MB-231-GFP (Cell Biolabs AKR-201) cell lines were then resuspended at 5000 cells/ml and 2500 cells/ml, respectively, in medium containing 10% FBS (Seradigm 1500), 1% NEAA (Thermo Fisher 11140), 1% pen-strep (Thermo Fisher 15140) in DMEM (Thermo Fisher 10569). After aspirating fibronectin, 50 ul of this cell suspension (totaling ~ 250 and 125 cells of the two cell lines, respectively) was then added to the bottom of each beat-plate well.

For glass-slide reaction chambers, 85 ul of growth medium (without cells) was added, and parafilm was used to cover the top of the reaction chamber assemblage. Holes were cut in the middle of cell culture wells (the four interior wells in Fig S1B). 10 ul pipette tips were then cut (S1C) and cell suspension was added from the wide end so that it traveled to the narrow end, and was held in place by capillary action. Parafilm was then added to wide end to create suction that would hold the cell suspension in place after the pipette tip was placed into growth medium. Pipette tips containing cell suspension and covered by parafilm were then placed vertically into the slide reaction chambers, and cells were allowed to settle.

Cells in all reaction chambers were then cultured 36-48 hours.

#### 1.4 In situ preparation

After culturing, growth medium was removed and cells were washed once with 1x PBS (prepared from Thermo Fisher AM9625). Cells were fixed in 4% formaldehyde (prepared from Thermo Fisher 28906) in 1× PBS for 15 minutes at room temperature. Formaldehyde solution was aspirated and replaced by 3 × PBS, and left for 10 minutes. Samples were washed twice for 10 minutes in 1 × PBS, and then permeabilized with a solution of 0.25% Triton X-100 (Sigma 93443) in 1× PBS for 10 minutes. Samples were then washed twice in 1× PBS, treated with 0. 1 N HCl (VWR BJ318965) for 2-3 minutes and then washed an additional three times in 1 × PBS. Samples were then kept at 4 C during preparation of the reverse transcription reaction.

Immediately before reverse transcription, samples were rinsed once in ddH_2_O. After aspiration, reverse transcription mixes were added containing 400 uM dNTP (Qiagen N2050L), Superase-In (Thermo Fisher AM2696) at 1 U/ul, Superscript III (Thermo Fisher 18080) at 10 U/ul, 1× Superscript III buffer, and 4 uM DTT. RT ultramers containing UMI’s (Table S1 for 4-plex, Table S4 for 24-plex) were included at 850 nM (for Samples 1-2) or 100 nM (for all others) each. These reactions were then incubated 60 C for the 3 minutes, followed by 42 C for 1 hour, and then held at 4 C. After aspiration, samples were washed three times in 1 × PBS, and kept at 4 C in the final wash overnight. Samples were rinsed with ddH_2_O, and after aspiration, 40 ul of an enzymatic digestion mix was added including 1 × exonuclease-I buffer (NEB B0293S) and 1.4 U/ul exonuclease I (NEB M0293). Reactions were incubated at 37 C for 40 minutes, and then washed three times in 1 × PBS.

**Figure S1:**
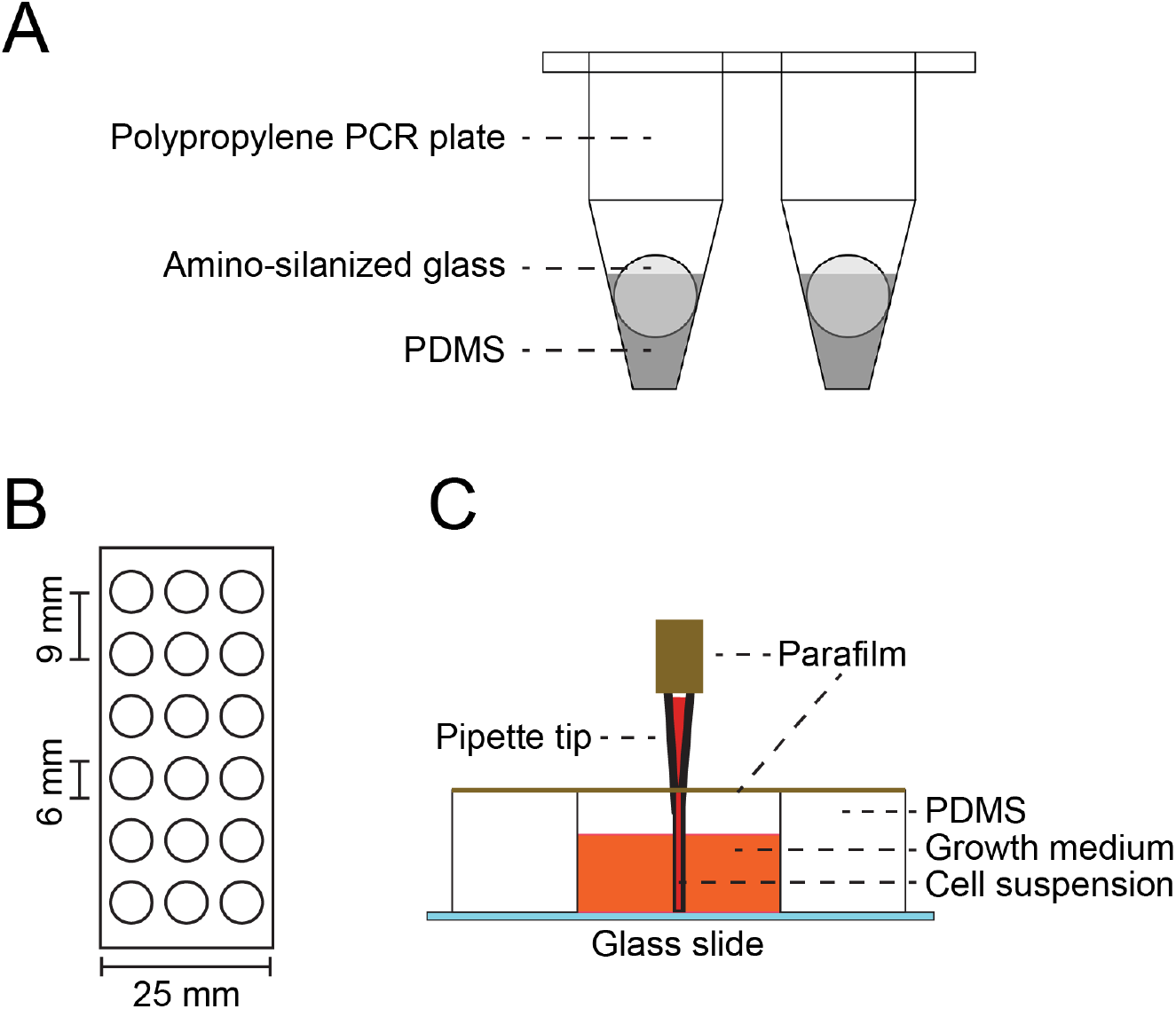
DNA microscopy reaction chambers, related to Fig. 1. (A) Bead-plate reaction chambers for the DNA microscopy on samples 1-2 and 4-5. Uncured PDMS is centrifuged to the bottom of polypropylene PCR plates. APTES-treated glass beads (coated with primary amines) are then added and spun into the uncured PDMS. The ensemble is then cured to generate a reaction chamber suitable for cell culture, multichannel pipetting, thermocycling, iterative enzymatic reacitons, and post-PCR containment. (B) PDMS cut used for glass-slide reaction chambers used to process sample 3. The interior four wells are used for cell plating, whereas the wells along the slide perimeter are used as reservoirs, containing 1× PBS. (C) Side view of glass slide reaction chamber when plasma-bonded to glass.

**Figure S2:**
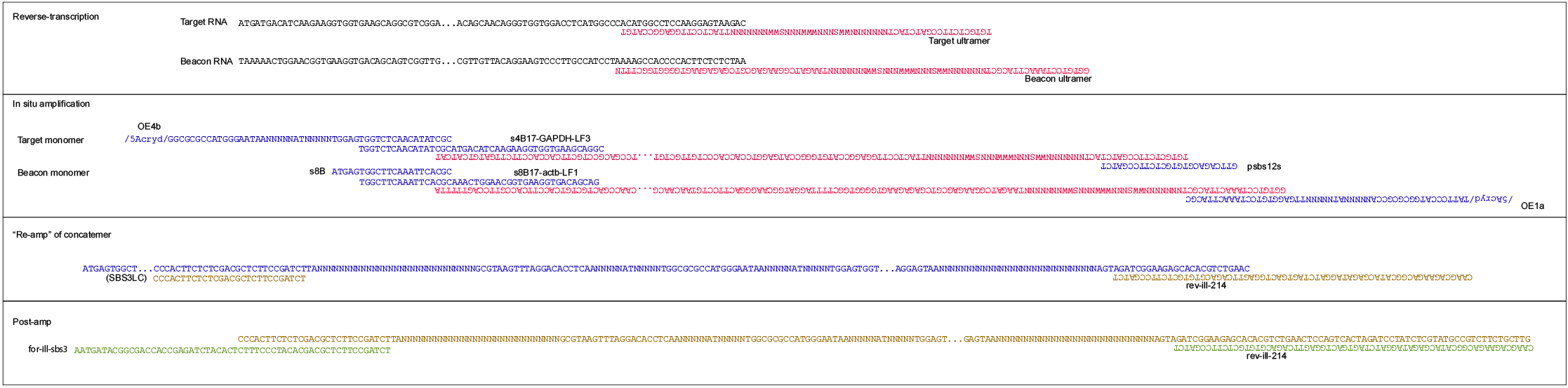
Assembly of the DNA microscopy amplicon in multiple steps, related to Fig. 1. The product achieved from the post-amplification step contains Illumina paired-end sequencing adapters.

Amplification mixes were prepared that included 400 nM each of primers OE1a and OE4b, 300 nM each of primers psbs12s (Lbs12s for 24-plex samples, Table S5) and s8B, 30 nM each of LF-primers (Table S2, or for 24-plex amplification, 10 nM each of the sF-primers in Table S5), 1.6 mM MgCl_2_, 200 uM dNTP, 0.5 mg/ml BSA (NEB B9000S), 8% glycerol (Thermo Fisher 15514011), Platinum Taq DNA polymerase (Thermo Fisher 10966018), 1× Platinum Taq PCR buffer, a 4-arm acrylate PEG (Laysan Bio 4ARM-PEG-ACRL-10K) at 64 ug/ul, and a 2-arm thiol PEG (Laysan Bio SH-PEG-SH-3400) at 44 ug/ul. Solutions were prepared in two parts, one containing the 2-arm thiol PEG, BSA, and glycerol, and one containing all other components. Following an additional sample rinse with ddH_2_O and aspiration, these two distinct components were mixed by pipetting and immediately added in 20 ul volumes (as a combined mixture) to the sample to allow for a 10.8% w/v hydrogel to polymerize for 1 hour at room temperature. This hydrogel would slow diffusion during the amplification reaction *(18)*.

Samples were then thermo-cycled at 95 C 2 min, 10×(95 C 30 s, 68 C 1 min), 2×(95 C 30 s, 55 C 30 s, 68 C 1 min), 16×(95 C 30 s, 60 C 30 s, 68 C 1 min), 68 C 1 min, 4 C. For 24-plex samples, samples were instead thermocycled 95 C 2 min, 1 × (95 C 30 s, 55 C 30 s, 68 C 1 min), 10 × (95 C 30 s, 68 C 1 min), 1 × (95 C 30 s, 55 C 30 s, 68 C 1 min), 16 × (95 C 30 s, 60 C 30 s, 68 C 1 min), 68 C 1 min, 4 C. The initial sets of 10 cycles at high temperature in these programs were designed to prime only one end of the cDNA amplicon. This would thereby confine initial amplification to increasing molecule copy numbers linearly with time, rather than exponentially. It would thereby minimize the effect of potentially stochastic amplification start-times.

Following in situ amplification, samples were stored at -20 C until further use.

#### 1.5 Library preparation

Frozen amplified samples were allowed to thaw on ice. A PEG-dissolution mix containing 460 mM potassium hydroxide (VWR BJ319376), 100 mM EDTA (Sigma 03690), and 40 mM DTT (Thermo Fisher P2325) was added directly on top of the hydrogel at 4 ul per sample while the sample was still on ice, and left for 2 hours at that temperature. Samples were then heated to 72 C 5 minutes, and mixed by pipetting 10 times. 4 ul of a neutralization solution made by combining 400 ul 1N HCl (Sigma H9892) and 600 ul 1M Tris-HCl pH 7.5 (TekNova T5075), adding this to the samples, and immediately mixing the solution again by pipetting. 11.1 ul of a proteinase mix was then added that contained 0.35% Tween 20 (Sigma P9416) and 0.35 mg/ml proteinase K (NEB P8107) in 10 mM Tris-HCl pH 8 (TekNova T1173). After mixing the samples by pipetting, incubation was performed at 50 C for 25 minutes.

55 ul of 10 mM Tris-HCl pH 8 was then added to each sample, and mixed by pipetting. 85 ul of the mixture was transferred to a new PCR plate, and 0.65 × volumes of Ampure XP beads (Beckman Coulter A63881) were added, mixed by pipetting, and left to incubate at room temperature 10 minutes. After twice washing with 70% ethanol, DNA was eluted into 35 ul 10 mM Tris-HCl pH 8. Product was then diluted 1:2 into a PCR reaction containing final con-centrations of 300 nM SBS3LC primer, 300 nM rev-ill-214 primer, 3.3 uM each of 10T-0E-P and 10T-0Ec-P interference primers (following on the strategy employed in Turchaninova et al (14) to prevent new concatemers from forming), 0.02 U/ul Platinum Taq HiFi DNA polymerase (Thermo Fisher 11304029), 1× Platinum HiFi Buffer, 1.5 mM MgS0_4_, and 200 uM dNTP. Reactions were thermo-cycled 95 C 2 min, 20×(95 C 30 s, 68 C 2 min), 4 C.

**Table S1:**
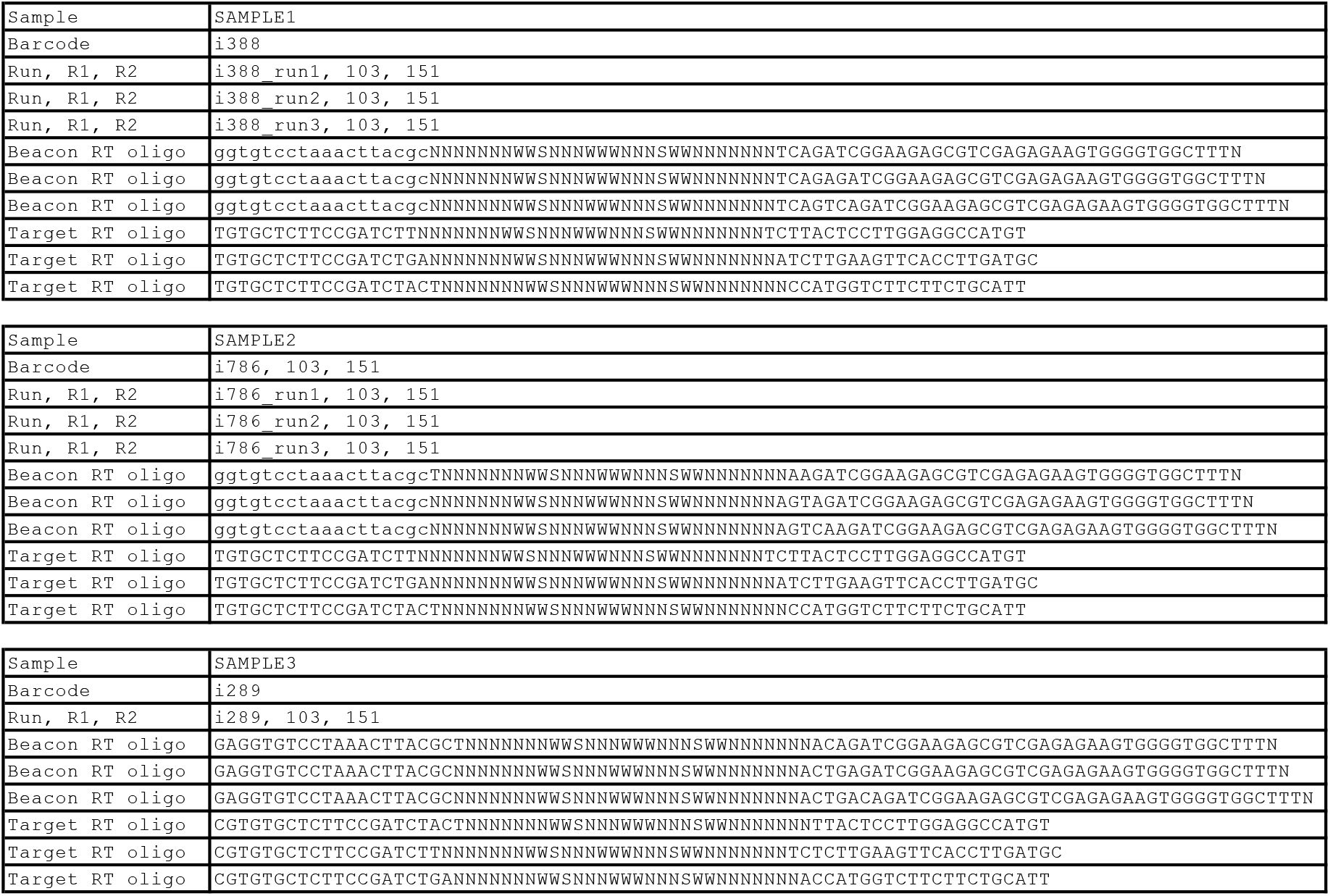
Oligonucleotides used for each sample during 4-plex (ACTB, GAPDH, GFP, RFP) reverse transcription, related to Figs. 2, 4–6, and S5. Lower case nucleotides indicate sequence areas during read parsing for which a 6% error rate is accepted, whereas upper case nucleotides afford zero error tolerance. Read-lengths labeled R1 (beginning at the 3’ end of the beacon UMI) and R2 (beginning at the 5’ end of the target UMI) are shown. All reverse transcription oligonucleotides were obtained as ultramers from IDT Inc.

**Table S2:**
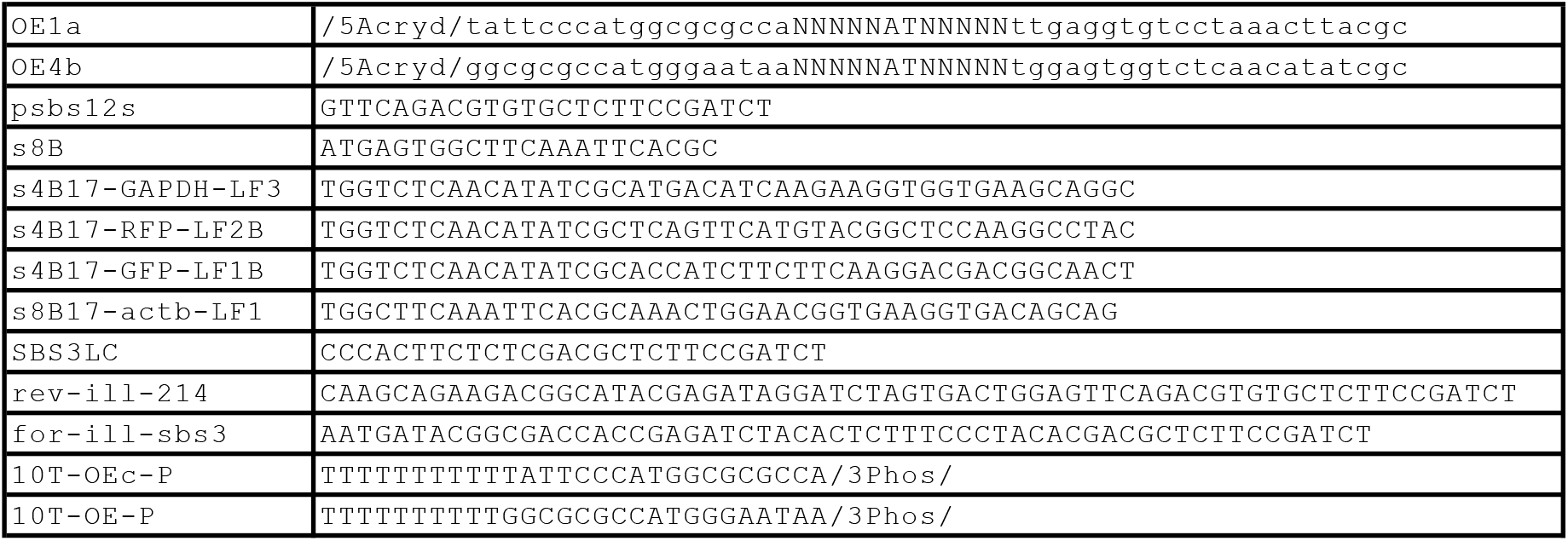
Oligonucleotides used for all samples during amplification and library preparation, related to Figs. 4–6 and Fig. S5. Lower case nucleotides indicate sequence areas during read parsing for which a 6% error rate is accepted, whereas upper case nucleotides afford zero error tolerance. 5’-acrydite modified oligonucleotides were HPLC-purified by the manufacturer.

Reaction products were Ampure XP-purified just as before, with 0.65 × volumes of Ampure XP beads added, and eluted into 40 ul 10 mM Tris-HCl pH 8. As part of a final sequence-barcoding step, 10 ul of sample eluent was added to a reaction containing 300 nM for-ill-sbs3, 300 nM rev-ill-X (with a sample-specific barcode where indicated on the sequence), 0.02 U/ul Platinum Taq HiFi DNA polymerase, 1× Platinum HiFi Buffer, 2 mM MgSO_4_, and 200 uM dNTP. Reactions were then thermo-cycled 95 C 2 min, 5×(95 C 30 s, 58 C 30 s, 68 C 2 min), and 1-5 × (95 C 30 s, 68 C 2 min), 4 C in order to obtain sufficient DNA library for sequencing.

#### 1.6 Sequencing

Following a final Ampure XP purification as above, with 0.7 × volumes of Ampure XP beads added, NGS libraries were sequenced on an Illumina NextSeq 550 instrument using manufacturer-standardized protocols for paired-end sequencing. Sequenced reads were de-multiplexed using the Illumina bcl2fastq pipeline using the 8nt sequence-barcode included 5’-adjacent to the SBS12 adapter 5’-GTGACTGGAGTTCAGACGTGTGCTCTTCCGATCT-3’ (**Fig. S2**, Table S2). Paired-end reads were sequenced from the SBS3 sequencing primer 5’-ACACTCTTTCCCTACACGACGCTCTTCCGATCT-3’ to 103 bp, and from the SBS12 sequencing primer to either 57 bp or 151 bp, as indicated in Tables S1 and S4, depending on whether full amplicon sequences were intended to be captured (instead of simply a minimum number of identifying bases).

#### 1.7 Additional considerations

##### 1.7.1 UMI/UEI design

The number of N’s to use in a UMI/UEI will depend on the expected diversity of molecules and/or events being tagged. Assuming an upper-bound for this diversity is known, the question reduces to the so-called “birthday-problem”. Given a UMI/UEI length ℓ (with each of ℓ bases having all 4 base possibilities), the probability that two randomly-drawn UMIs/UEIs will match (assuming uniform base-distributions) is *P*_0_(ℓ) = 4^−ℓ^. Similarly, the probability that there will be another UMI/UEI within 1 bp is

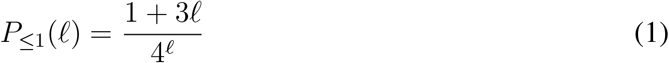

because there is 1 way for a randomly drawn sequence to be precisely the sequence of a previously drawn sequence, and 3ℓ ways for it to be the same except for exactly 1 mismatch. The probability that no two UMIs/UEIs out of *N* will overlap in this way is

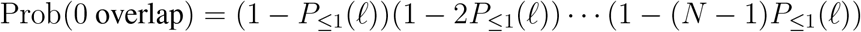

Define *N*_crit_(ℓ) through the relation

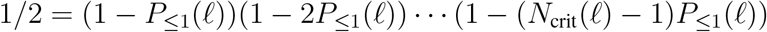

Then *N*_crit_(ℓ) is the maximum diversity of templates beyond which it becomes likely that at least 1 pair of UMI/UEI sequences will be within 1 bp of one another. For UMI/UEI sequences in which there are ℓ_4_ bases that are randomly selected across all 4 nucleotides and ℓ_2_ bases that are randomly selected across 2, we can re-write equation 1:

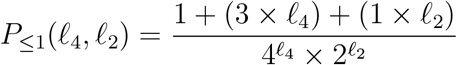

Since the UMIs used in our experiments (Table S1, Table S4) have ℓ_4_ = 20 and ℓ_2_ = 9, this gives us N_crit_ = 3.3 × 10^6^ for each beacon- and target-UMI data set presented here.

Note that for UEIs, the picture is far simpler. Because a UEI brings together exactly two UMIs, two UEIs that are grouped together will get one vote (assigned via plurality). Therefore, the less abundant indistinguishable UEI will simply be ignored. From here we can see that we can bring UEI diversity far closer to the upper limit of that which is physically possible (4-, or in our case ~ 10^12^) without substantial problems.

**Table S3:** Read and UMI/UEI counts for all samples, related to Figs. 2, 4–6, S5. Columns labeled “> 1 read” denote status of UMIs/UEIs identified by independent clustering, whereas columns labeled “MLE” denote those that made it into the final UEI matrices for image inference. “UEI/Beacon UMI” and “UEI/Target UMI” denote ratios, important for analyzing data quality in the context of resolution defined in equation 11. Asterisks denote 24-plex samples, for which amplicon-filtering was done after UMI/UEI clustering, and therefore those discarded on this basis were not counted during initial read parsing.

| Sample | Total sequenced | Fraction retained | Beacon UMIs (>1 read) | Target UMIs (>1 read) | UEIs (>1 read) |
| --- | --- | --- | --- | --- | --- |
| 1 | 26832238 | 0.76 | 892372 | 535216 | 5056844 |
| 2 | 21355785 | 0.76 | 898039 | 488100 | 3915689 |
| 3 | 34846809 | 0.60 | 63857 | 31960 | 732428 |
| 4 | 44032216 | 0.82* | 402575 | 109482 | 1291422 |
| 5 | 42669570 | 0.85* | 152875 | 48577 | 457148 |

| Sample | Beacon UMIs (MLE) | Target UMIs (MLE) | UEIs (MLE) | UEI/Beacon UMI (MLE) | UEI/Target UMI (MLE) |
| --- | --- | --- | --- | --- | --- |
| 1 | 413574 | 268752 | 4149473 | 10.0 | 15.4 |
| 2 | 329848 | 215285 | 2989182 | 9.1 | 13.9 |
| 3 | 38307 | 21192 | 695307 | 18.2 | 32.8 |
| 4 | 146653 | 48354 | 757502 | 5.2 | 15.7 |
| 5 | 52359 | 19951 | 274540 | 5.2 | 13.8 |

Note furthermore that even for UMIs, things get easier if the target sequences are used to separate out UMIs (this is not done for any data-set presented here). For a set of target sequence frequencies {*p*_1_,*p*_2_,…,*p_s_*} (normalized to sum to one) of S distinct sequence-types labeled by UMIs, the probability that two randomly selected sequences will be the same is 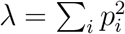. This measure, also known as Simpson’s diversity index, affects the calculation above by multiplying *P*_≤1_. The more diverse and distributed the population of sequences, the smaller the product *λP*_≤1_ and the larger the value of *N*_crit_(ℓ).

##### 1.7.2 Remark on reagent quality control

Although reagent quality control is crucial for every protocol, PEG reagents used for the in situ PCR step are especially sensitive to variation. Basic precautions that must be taken include desiccation with Drierite (Sigma 238961) or a similar agent in a sealed bag at -20 C. Lot-to-lot variation must be controlled by keeping a careful log of the lots used for each experiment. We found that in general this variation could be pre-checked by performing routine bulk PCR’s within the hydrogel, and comparing the results on a gel. UV/Vis comparison may also be used as a way to compare inorganic salt content that may have carried over from manufacture.

### 2 Analysis

#### 2.1 Read parsing

Reads were parsed by first gating out those with a mean quality score of less than 30 on either the forward or reverse read. Forward and reverse reads were then checked for inclusion of primer and stagger sequences, depending on the primers used and indicated in Tables S1 and S4. Capitalized bases were used to indicate those base positions intolerant of a single mismatch, whereas consecutive stretches of lower-case bases were used to indicate pieces of sequence that were permitted to contain mismatches up to the indicated maximum of 0.06 as a fraction of total. For 4-plex data, 5 bases after gene-specific primer sequences were used to gate reads in order to remove unrecognized gene inserts. Read counts and fractions retained for each data set are shown in Table S3.

#### 2.2 UMI/UEI clustering

In order to identify UEI and UMI sequences in a way that would make efficient use of the data available and in a manner specifically accommodating to long-tailed distributions of PCR error, we developed a simple clustering algorithm, which we here on refer to as EASL, or Extended Abundance Single-Linkage. EASL relies on single-mismatch alignments alone to identify clouds of erroneous sequences that decay in density the further in sequence-space they exist from an abundant, putatively correct, original sequence.

EASL clustering (**Fig. S3A**) initiates by grouping every UMI/UEI (from each read location separately, so that it disregards the rest of that read) within a data set by perfect identity. The abundance (by read-count) is assigned to each UMI/UEI sequence. Each pair of UMIs/UEIs is compared by un-gapped alignment. This may be performed by local similarity hashing in a way that permits full pairwise comparison in *O*(*NL*^2^) time, where N is the number of unique UMIs/UEIs, and L is the length of the UMI/UEI sequence. In brief, this may be achieved by generating L hash-tables/dictionaries of UMI/UEI sequences, where each of these dictionaries has a specific sequence-position removed. For each dictionary, the full collection of L — 1 length UMI/UEI sequences generated by removing that corresponding position are added to the dictionary. Those found grouped together will be those sets related by a single base mismatch.

EASL clustering then proceeds as follows (**Fig. S3A**). UMI/UEI *i* **directionally** links to UMI/UEI *j* if and only if the read-abundance of UMI/UEI i is greater than or equal to the read-abundance of UMI/UEI *j*. Read number densities (RNDs) are calculated for each UMI/UEI sequence by summing read-abundances belonging both to the sequence itself and to all sequences (of equal or lower abundance) it links to. Each UMI/UEI data set is then independently sorted by decreasing RND, and accordingly clustered independently, as follows.

The UMI/UEI with the largest RND initiates clustering as the first cluster-seed. All UMIs/UEIs to which this seed links by the aforementioned criterion are accepted into its cluster. The algorithm then proceeds to the UMI/UEI with the next largest RND that has not already been assigned a cluster. This UMI/UEI becomes a new cluster seed and all UMIs/UEIs not yet assigned to a prior cluster are accepted to that belonging to the new seed. This process proceeds among all un-assigned UMIs/UEIs down the RND-sorted list. When no un-assigned UMIs/UEIs remain, the algorithm terminates.

**Figure S3:**
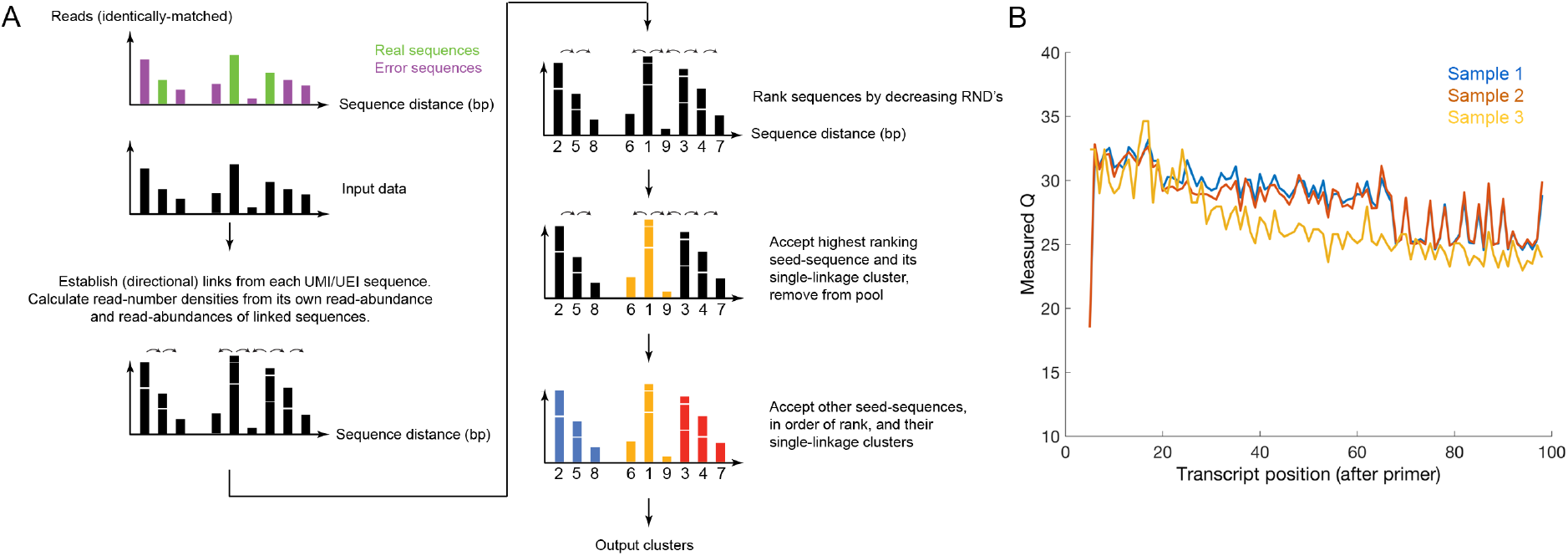
Sequence-error handling at the level of UMIs, UEIs, and transcript inserts, related to Fig. 1. (A) An illustration of the EASL clustering method for UMI and UEI sequence clustering in log-linear time. (B) Quality score (– 10 log_10_ Prob(incorrect)) dependence on position for target amplicons belonging to GFP and RFP *after the annealing primers* in Table S1. Samples 1, 2, and 3 (blue, red, and yellow, respectively) were sequenced out to ~ 100 bp past the annealing primer site. and they are therefore shown here. Plot begins ~ 5 bp into the transcript, since the first 5 bp were used during initial read-filtering.

#### 2.3 UEI-UMI pairing

After clustering, UMIs were accepted to further analysis if they associated with at least two reads (UMIs and UEIs remaining after this preliminary filtering step are represented by the upper-curves in the rarefaction plots in Figs. 3–4 and S5). UEI clusters were then matched with the beacon UMI/target UMI cluster-pair to which they were found to associate with the most reads in the original data. This filtering was intended to remove incorrect associations between beacon UMIs and target UMIs caused by, among other things, PCR chimera formation during downstream library preparation steps (we reasoned that the original UMI-UEI-UMI pairings would have a head-start before late-stage amplification – once a new and incorrect UMI-UEI-UMI pairing would form, it would begin amplifying from a quantity of 1 molecule later in downstream amplification).

The resulting “consensus” UMI-UEI-UMI pairings were then iteratively filtered by eliminating UMIs associating with fewer than 2 UEIs. After the initial set of UMIs were removed on account of having too few UEIs, the UEIs they associated with were removed, the matrix was re-filtered to exclude UMIs that no longer had at least 2 UEIs, and so on, until no remaining UMIs could be found associated with fewer than 2 UEIs. The resulting pruned data set is shown in the lower-curves of the rarefaction plots in Figs. 3–4 and S5.

The target sequences grouped by UMI-clustering further allowed errors owing to PCR and sequencing to be suppressed. Aggregating the votes at every position for the reads grouped under a particular target UMI gave quality scores (−10 log_10_ Prob(incorrect)) that mostly stuck to Q=30 until 60-70 bp after the end of the primer, after which Q hovered between 25-30 until it ended nearly 100 bp into the transcript (Fig. S3B).

#### 2.4 Image inference

##### 2.4.1 Formalization

Consider the evolving concentration distribution of products of a single UMI with index *i*, centered at position 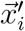, during a DNA microscopy reaction. This can be modeled as isotropic diffusion using the Gaussian profile for concentration at position 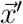 at time *t*:

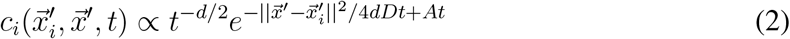

where *d* is the dimensionality (of physical space), *D* is the diffusion constant, and *A* = log 2/Δ*t* where Δ*t* is the time-scale of a PCR cycle. The rate of UEI/concatemer formation between UMIs *i* and *j* with the same diffusion constant will then be the volume-integral

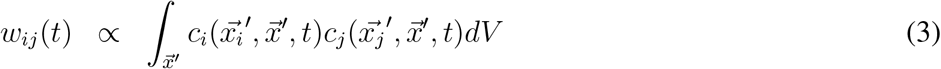

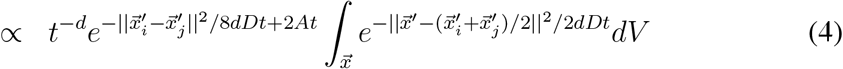

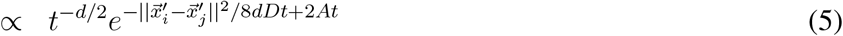

Note that although the UEI formation rate is time-dependent – and that therefore the total observed reaction rate is in fact a sum of functions above from each PCR cycle – provided amplification happens quickly, prior time-dependence to some final reaction time *τ* will be swamped out by the reaction rate at that time *τ*. Therefore, for the sake of simplicity, we will drop the time dependence from our probability model, and say that UMIs *i* and *j* located at *t* = 0 at positions 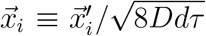 and 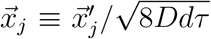, respectively, will have an expected cumulative reaction rate of

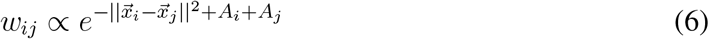

where *A_i_* and *A_j_*, are amplification “biases”, ie the cumulative effective amplitudes of the UMIs’ diffusion profiles. The length scale above is denoted

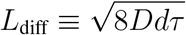

in the main text.

**Figure S4:**
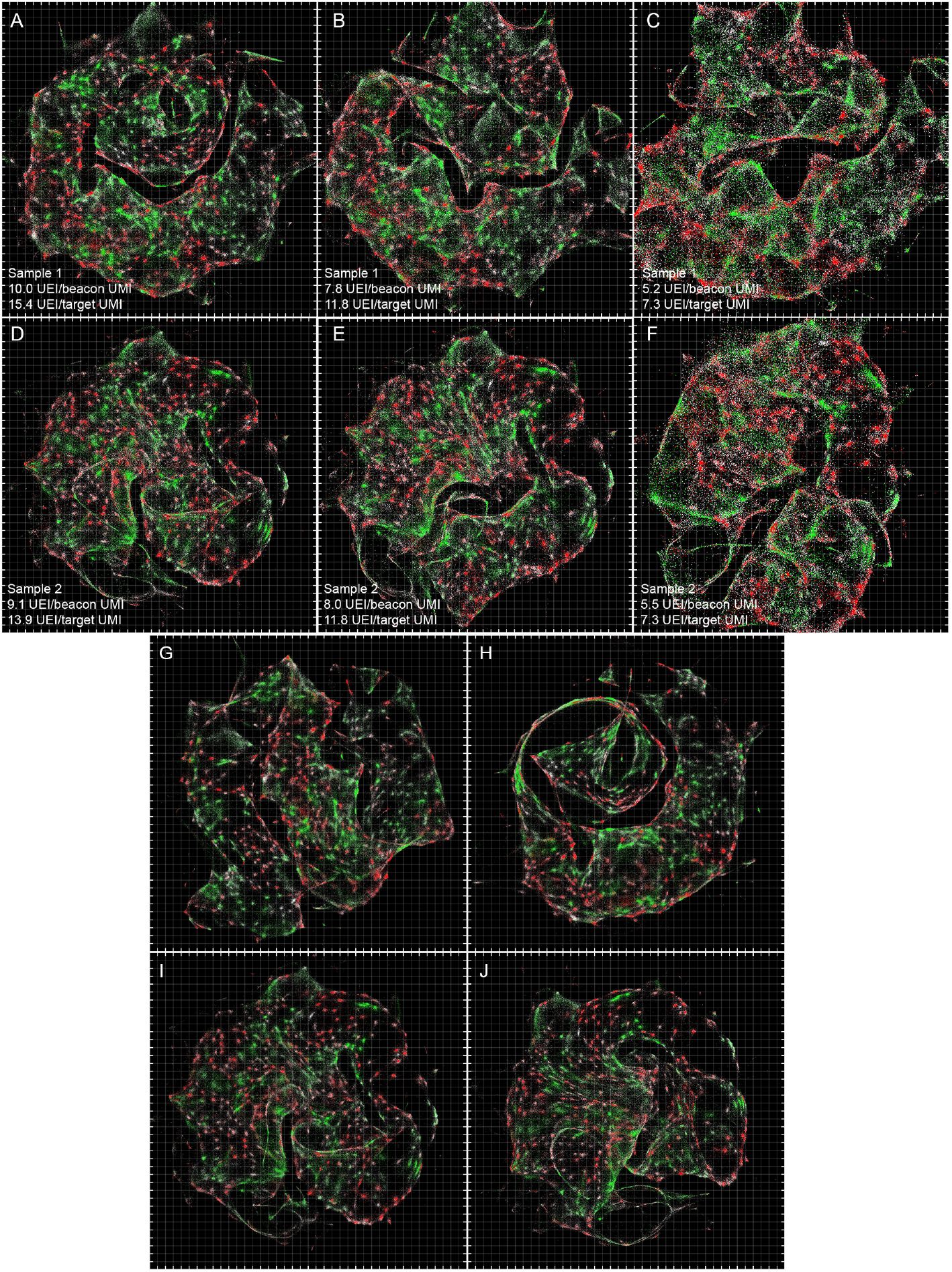
Data down-sampling and re-parameterization of sMLE inference, related to Fig. 5. Downsampling of samples 1 and 2 at various read-depths (A-C and D-F, respectively). A and D correspond to all-inclusive data-sets (20273379 retained reads for sample 1, 16248577 retained reads for sample 2). B and E correspond to 12800000 sub-sampled reads and C and F correspond to 6400000 sub-sampled reads. sMLE-initialization to 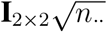 for samples 1 (G) and 2 (I), instead of initialization to just the identity matrix **I**_2×2_, the conditions used in panels A and F. sMLE inference stopped at 50 (instead of 100) eigenvectors for samples 1 (H) and 2 (J).

The probability of observing UEI counts {*n_ij_*} for each UMI-pair 〈*i,j*〉 is then the multinomial expression

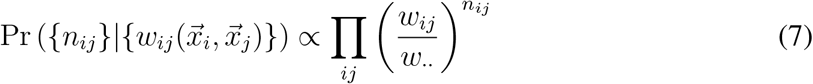

where dots “·” represent index summation (so that *w*·· ≡ Σ*_ij_ w_ij_*). From this, we can write the log-likelihood

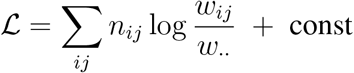

and, relying on the functional form from equation 6, its gradient with respect to UMI position 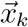

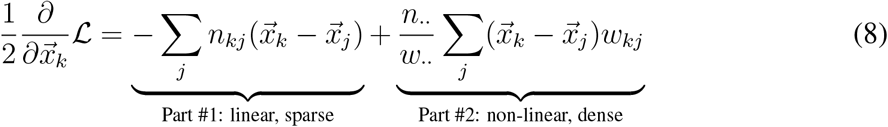

A solution to the above occurs when the gradient is zero for every 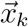, and where contributions from equation 8 parts #1 and #2 balance. Parts #1 and #2 of equation 8 differ in several important ways.

The first of these differences is the role they play: part #1 dictates how to **center** each UMI relative to one another, and attracts all UEI-associated UMIs together, whereas part #2 regulates how to **separate** them, by repelling all UMIs from all other UMIs at the strengths dictated by the intrinsic lengthscale of the function *w_ij_*. The second difference is the ease of calculation: part #1 involves summing only the UEIs observed in the experiment, the summation is sparse in the same way our observations are sparse; part #2 meanwhile makes no distinction between UEIs that are observed and UEIs that are not, and requires the summation over all UEIs that are *possible.*

Third and finally, the two expressions differ in the length scales over which they operate. Optimization over small length scales containing minimal point density variation will make part #2 of equation 8 approach zero, since it involves the summation of repulsive forces pointed in all directions. In these circumstances, part #1 will dominate. However, over long distances in which large-scale point densities may vary, part #2 will contribute heavily.

##### 2.4.2 Local linearization of the image inference problem

We can write part #1 of equation 8 as 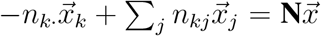, where 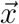 is now a solution to all UMI positions simultaneously, and where we’ve defined the zero row-sum UEI matrix, or what in the main text is referred to as the UEI Graph Laplacian:

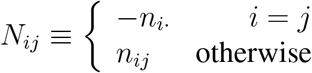

Note that **N** is a sparse square matrix (all UMIs × all UMIs), re-written from the rectangular form in Fig. 1E, which has exclusively beacon UMIs as rows and exclusively target UMIs as columns. **N** will always have a “trivial” eigenvector of all 1’s (with eigenvalue 0) that solves the equation by making all positions equal. Solving equation 8 part #1, by obtaining the nontrivial solution nearest to 0 means setting 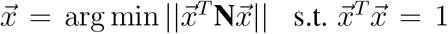, and amounts to maximizing the numerator of the multinomial probability in equation 7. We will write the solution to this eigenvalue problem in a row-normalized form

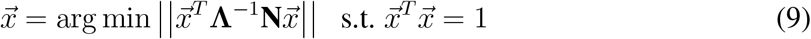

where we use the diagonal matrix

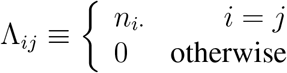

to equalize contributions to the gradient by each UMI.

Because the solution to the maximum likelihood problem is only linear locally, we need a way to zoom in on local portions of the data in order to use it. We can do this by applying simple and approximate graph cut/spectral partitioning algorithms previously described *(15).* Specifically, we take the symmetric normalized form of the UEI Graph Laplacian, Λ^−1/2^NΛ^−1/2^, and find its second smallest-in-magnitude (after the trivial solution) eigenvalue/eigenvector pair. We then perform a sweep of possible cuts within that eigenvector to minimize the conductance between the resulting UMI sub-sets *A* and *B*: *N*(*A, B*)/ min(*N*(*A*), *N*(*B*)), where *N*(*A, B*) is the number of UEIs associating UMI sub-sets *A* and *B*, and *N*(*A*) and *N*(*B*) are the total number of UEIs belonging to those two UMI sub-sets, respectively.

Minimizing this conductance value allows for the “sparse cut” described in the main text. By iteratively cutting the matrix, re-forming the matrix Λ^−1/2^NΛ^−1/2^, and continuing until the minimum available conductance-cut is above a threshold, we can obtain local linear data subsets depicted in Fig. 2E-H.

Setting the threshold higher provides for more extensive cutting, and results in the cell segmentation shown in Fig. 6A-B. The accompanying binomial p-values in Fig. 6C-D present, for putative cells with ≥ 50 UMIs and at least one ACTB and one GAPDH transcript:

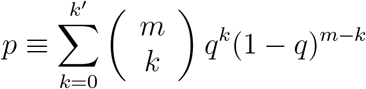

where *k*’ is the number of UMIs belonging to the minority transgene (GFP or RFP) within that putative cell, *m* is the total number of transgene UMIs observed, and *q* is the frequency of the minority transcript within the entire data set. It therefore describes the probability of observing the cell as-is under a random partitioning hypothesis.

##### 2.4.3 Global likelihood maximization

Moving to larger length scales means dealing with part #2 of equation 8. In order to handle the large-scale summation of every pair of UMIs (otherwise prohibitive due to its quadratic scaling), we adapted the Fast Gauss Transform (17) that allowed calculation of this sum with bounded error in linear time. Error bounds were parametrized as the maximum possible fraction of the calculation of the weight-sum *w_k_*. for each UMI. This was set to 30%, which sufficed to constrain actual error levels to orders of magnitude smaller (Fig. S6).

**Figure S5:**
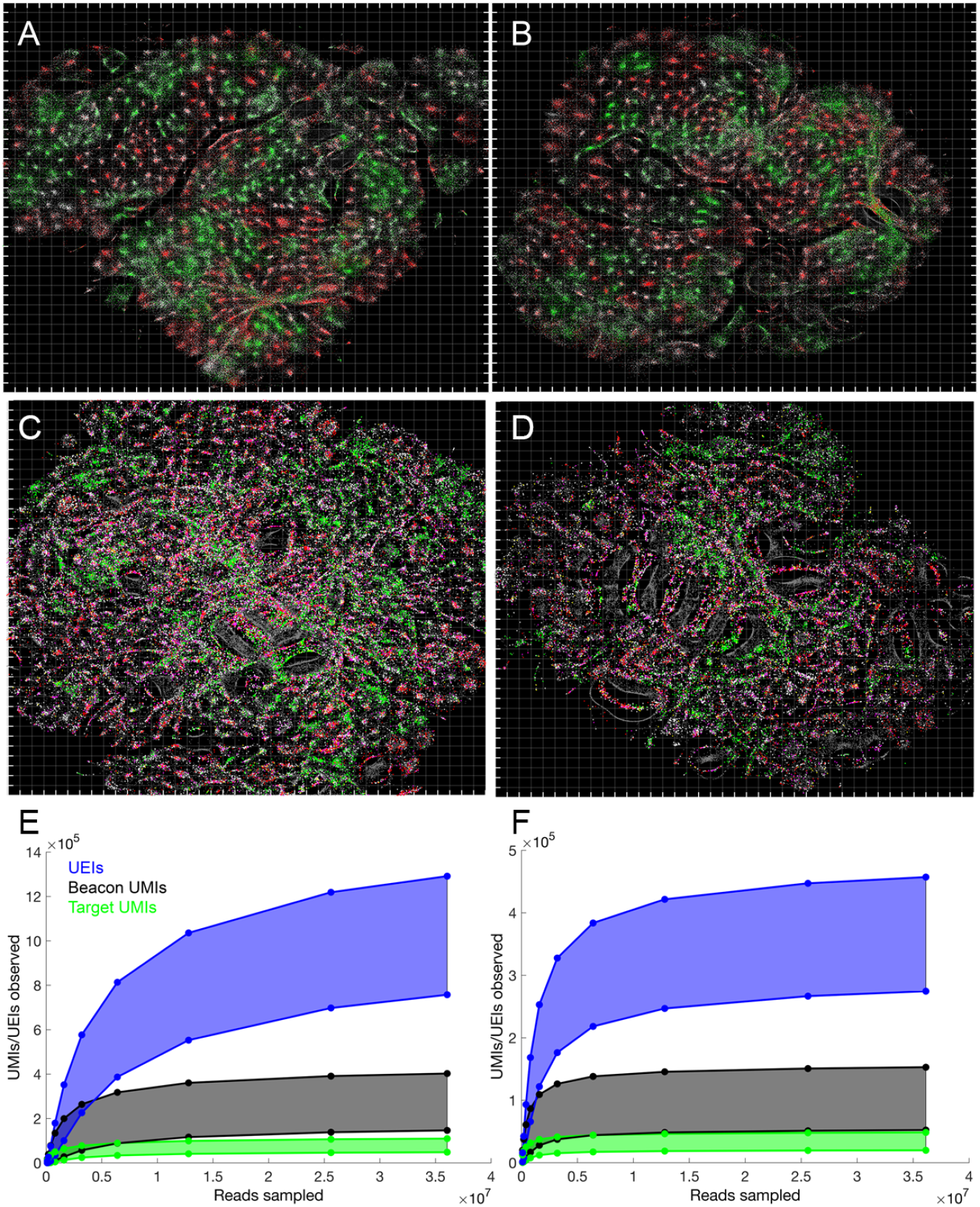
Global point-MLE solutions for 4-plex and 24-plex gene targeting, related to Figs. 5–6. Point-MLE solutions for 4-plex samples 1 (A) and 2 (B), in which each UMI is optimized independently. Grid-lines are used to denote spacings of *L_diff_* (or *Δx* = 1.0 in equation 8, used for image inference). For 4-plex data, grey = ACTB/beacon, white = GAPDH, green = GFP, and red = RFP. (C-D) Point-MLE solutions for samples 4 and 5, respectively, with 24-plex targeting performed on sub-sets of genes known enriched in BT-549 and MDA-MB-231 cell lines (Tables S4 and S5). All targeted genes were found at non-zero frequencies except for GRIN2D, MEA1, FAM170B, and C11ORF44. Gene colorings are the same as in A-B, but included are genes previously found enriched in the MDA-MB-231 (yellow), which here expresses GFP, and BT-549 (magenta), which here expresses RFP. Rarefaction plots for samples 4 (E) and 5 (F) are also shown, with top and bottom curves indicating data immediately following clustering and data entered into the UEI matrix for image inference, respectively.

**Figure S6:**
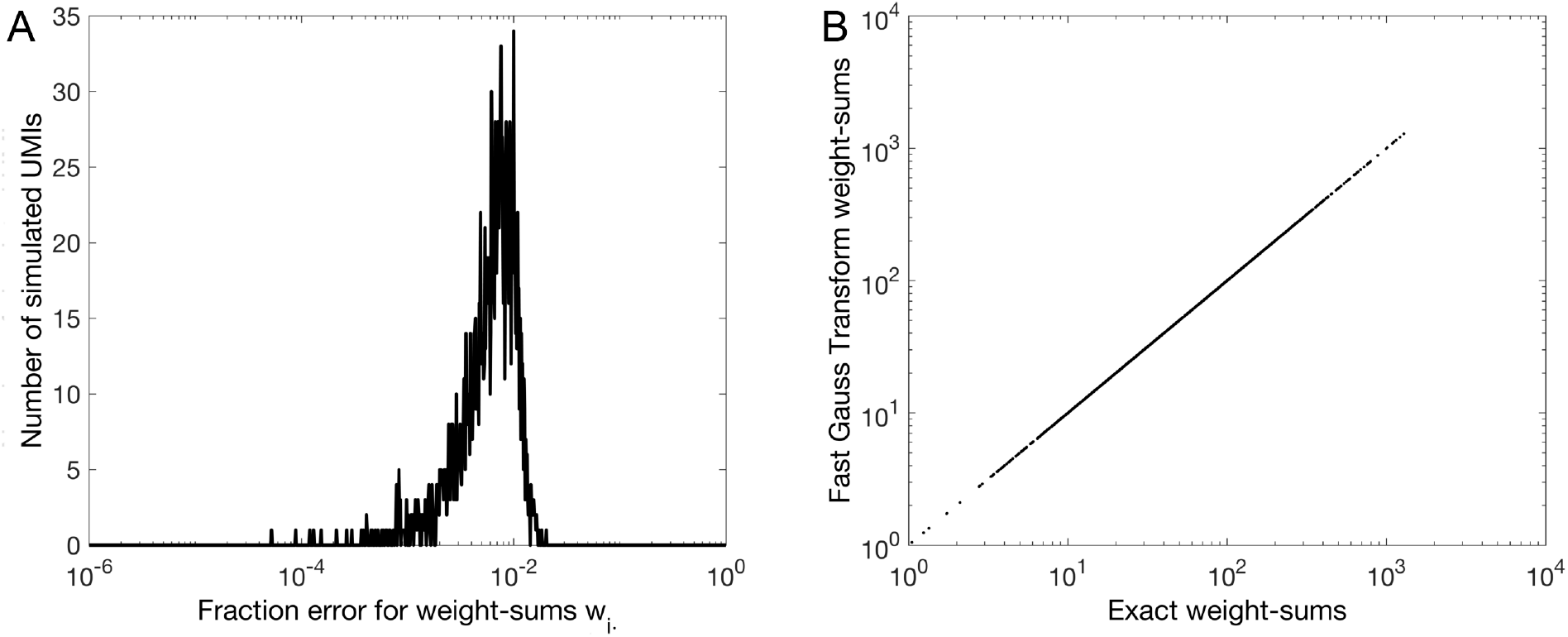
Simulated fractional error histogram (A) and correlogram (B) using an adapted Fast Gauss Transform, related to Fig. 3–6. The Fast Gauss Transform allows sums of *O*(*N*^2^) Gaussian interactions between point-sources (here, UMIs) to be calculated in *O*(*N*) time. The Fast Gauss Transform is applied above to the weight sums *w_i_*. = *Σ_j_ w_ij_* of 2000 simulated UMIs (1000 beacons and 1000 targets) normally distributed with *σ* = 10, and with amplitudes *A_i_*,*A_j_* (from equation 6) normally distributed with *σ* = 1. Maximum fractional error bound is set to 30% for weights *w_i_*.

##### 2.4.4 Point-MLE solution

The most straightforward way solve equation 8 is to randomly initialize the global solution with an “educated guess” of what the global solution might look like and perform a gradient ascent of the global likelihood function using each UMI position as an independent variable. Since the eigenvector solutions to equation 9 satisfy local constraints, they provide a logical starting-point to initialize this solution. To this end, we let the top 100 eigenvectors of the **full** data-set’s row-normalized UEI matrix Λ^−1^ N be columns in the matrix **Z**. The *d*-dimensional (with all samples here having *d* = 2) initial UMI positions 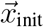 were then defined through the linear combination

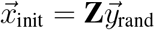

where 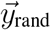 was a *d*-column matrix, with each row corresponding to a different eigenvector, and its elements being linear coefficients used to sum the columns of eigenvectors in **Z**. The elements of 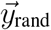 were normally distributed coefficients generated from 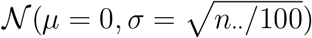). Amplitudes *A_i_* were set to log *n_i_*. (an approximation asserting that, on average, UMI density was uniform at the length scale of diffusion) and gradient ascent proceeded using the calculation from equation 8 (and applying the L-BFGS optimization method from the SciPy library). This point-MLE approach was applied in Fig. S5A-D.

These global solutions illustrate, however, the difficulty in capturing information on empty space when each point is being optimized independently. Clusters of points are unable to separate by more than the length-scale *L*_diff_ indicated by grid-lines (which is defined as the unit-less value of 1.0 in the physical model of equation 6).

##### 2.4.5 Spectral MLE (sMLE) solution

In order to capture more information on empty space than the point-MLE solution allows, we can expand on the local linear solutions previously described, and require our global solutions to *remain* linear combinations of the top eigenvectors of the full data-sets’s row-normalized UEI matrix Λ^−1^N. Again assembling these top eigenvectors as columns in the matrix **Z**, the global d-dimensional (with all samples here having *d* = 2) solution 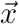 of size *M × d* for all *M* UMIs was then defined as the linear combination

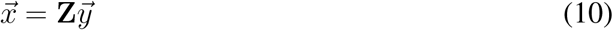

where, as before, 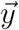 was a *d*-column matrix, with each row corresponding to a different eigenvector, and its elements being the linear coefficients used to sum the columns of eigenvectors in **Z**. Using equation 10 made 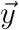 a low-dimensional variable set that we could optimize directly. The coefficients in 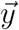 therefore dictated the UMI positions 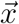 and the gradients of each UMI in 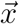 were then calculated using equation 8. These individual UMI gradients were then projected back onto the linear eigenspace defined by **Z**, allowing 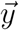 to be updated accordingly. Because eigenvectors in **Z** were not orthogonal, the back-projection of high-dimensional gradient 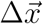 to low-dimensional gradient 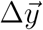 was defined (through equation 10) by 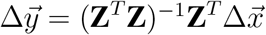.

We approached this low-dimensional optimization within the eigenspace **Z** in as incremental a way as possible. We called this incremental approach “spectral maximum likelihood estimation”, or sMLE. On iteration 1 of sMLE, the first 2 eigenvectors of matrix **Z** were taken in isolation (corresponding to the non-trivial eigenvalues with smallest magnitude), and performing a gradient-ascent optimization of equation 8 with their coefficients alone gave optimal coefficients for generating a solution 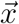 from a linear combination of these two eigenvectors alone. On iteration 2, the eigenvector with the next smallest-magnitude eigenvalue/eigenvector pair in **Z** was added to those allowed to contribute to the solution 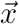, thereby adding to the number of optimizable coefficients in 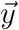. This larger vector 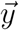 was then optimized for the now 3 eigenvectors. This was repeated until all top eigenvectors (numbering 100 in the presented data sets) were integrated into the linear combination defining solution 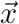.

The outputs are plotted in Figs. 4–6 and – for down-sampled read counts – in Fig. S4A-F. It should be noted that the two parameters that need to be fixed for this algorithm are the scaling factor that multiplies the initial 2 eigenvectors that seed the solution and the choice of total eigenvectors after which the algorithm terminates (Fig. S4G-J). Although alterations in the initial scaling factor result in isomorphic images (due to their using a common set of eigenvectors to construct a solution), certain manifold folding-defects can be mitigated by scaling factor choice. A comparison is made between initializing 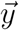 in equation 10 to the identity matrix **I** (applied to solutions in the main text and Figs. S4A, and S4D) versus initializing 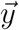 to 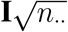, where *n*. is the total UEI count (applied to Figs. 4, S4G, and S4I). The total number of eigenvectors at which the sMLE algorithm terminates (shown at 50 in Figs. S4H and S4J, compared to 100 everywhere else) similarly alters manifold folding by freezing out certain degrees of freedom the solution can use to maximize the position likelihood function.

##### 2.4.6 Resolution and UEI count

The relationship between the uncertainty of a UMI’s position given its neighbors’ can be understood as the equivalent of the standard-error in a statistical average (namely, the standard-deviation divided by the square-root of the number of independent measurements). However we sketch it out explicitly here in the context of the solution-likelihood function in equation 7. If we assert that in regions where the local linear conditions previously discussed apply (gradients in point density at the diffusion length-scale *L*_diff_ are small), a solely varying UMI *k* has solution-likelihood at position 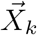 about some maximal likelihood 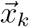

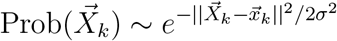

then we can simply calculate

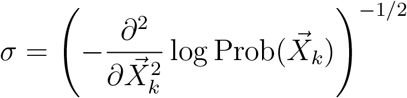

Since under the local linear conditions, the multinomial probability in equation 7 becomes a simple product of Gaussians, we get

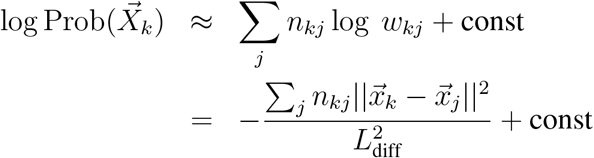

where we’ve retained the physical length *L*_diff_ in equation 6. From this we can finally write

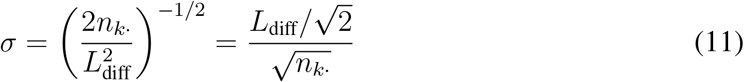

meaning a UMI’s positional uncertainty will shrink with the square root of the total number of UEIs with which it associates. This relationship is highlighted in Fig. 3B-C.

##### 2.4.7 Simulation

The efficacy of the sMLE algorithm was evaluated in a more controlled setting using simulated data exhibited in Fig. 3G.

Simulations proceeded as follows. For each UMI *i*, molecular-copy numbers *m_i_*(*t*) at amplification cycle *t* was initiated at *m_i_*(*t* = 0) = 1. For discrete *linear* amplification cycles *t* = 1, 2,…, *τ*_lin_, with *τ*_lin_ being the total linear-amplification cycle number, the total molecular-copy numbers were updated as

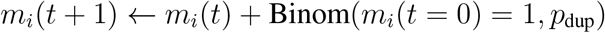

where 0 < *p*_dup_ ≤ 1 was the efficiency at which each template (UMI-tagged cDNA) molecule was copied. As in the experimental protocol, linear amplification was followed by exponential PCR amplification, in which molecular-copy numbers were updated as

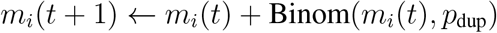

for *t* = *τ*_lin_ + 1, *τ*_lin_ + 2,…, *τ*_lin_ + *τ*_exp_. Meanwhile, during exponential PCR cycles *t* = *τ*_lin_ + 1, *τ*_lin_ + 2,…, *τ*_lin_ + *τ*_exp_ the expected rate of UEI formation *w_ij_*(*t*) between every beacon *i* and target *j* was calculated according to the previously derived equation 5

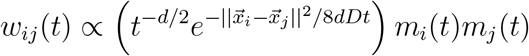

where the expectation values of the total molecular abundance of beacon UMI *i* and target UMI *j* are here explicit – *m_i_*(*t*) and *m,j* (*t*), respectively. For a given total *final* UEI count *N*, we then calculated an *expected* UEI count for time *t*

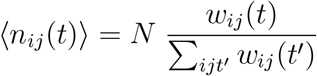

The number of actual UEI formation events for every triplet (*i, j,t*) were then assigned randomly using Poisson statistics

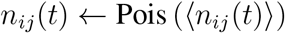

The *k*th UEI forming event generated by UMI-UMI pair (*i, j*), would then come into existence at its time-of-creation *t* with a molecular count *a_ijk_* (*t* = *t*’) = 1. That abundance would evolve in time until the end of the reaction *t* = *τ*_lin_ + *τ*_exp_ according to the iteration relation

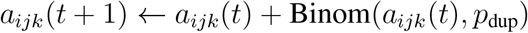

Each UEI’s final read-abundance, given an expected total read depth Ω, was then assigned

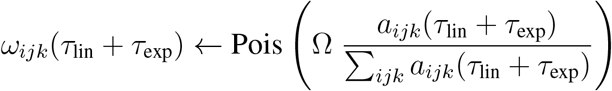

Image inference algorithms were then applied to this simulated data set in Fig. 3G. Here, freely diffusing products from 5000 beacons and 5000 targets, incorporating amplification stochasticity and sparse UEI sampling (50000 UEIs). “Original” coordinates are ground truth. UEIs in simulation are generated from an amplification reaction the same as in the experiment (see section *In situ preparation*), with 10 linear amplification cycles and 16 exponential amplification cycles. Amplification stochasticity was introduced by making each molecular duplication event 5% likely to not occur at all (*p*_dup_ = 0.95). Each cycle taking place over Δ*t* =1 with a *D* = 1 diffusion constant: both of these are in arbitrary units, with ground-truth positions normalized to the length scale 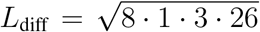 (the length scale from equation 5), since *D* =1, *d* =3, and *τ* = 26 (note that diffusion is still simulated in 3 dimensions, even though initial molecules are stationed in 2). Image inferences from simulations are re-scaled and registered (rotation/reflection) relative to ground-truth.

**Table S4:**
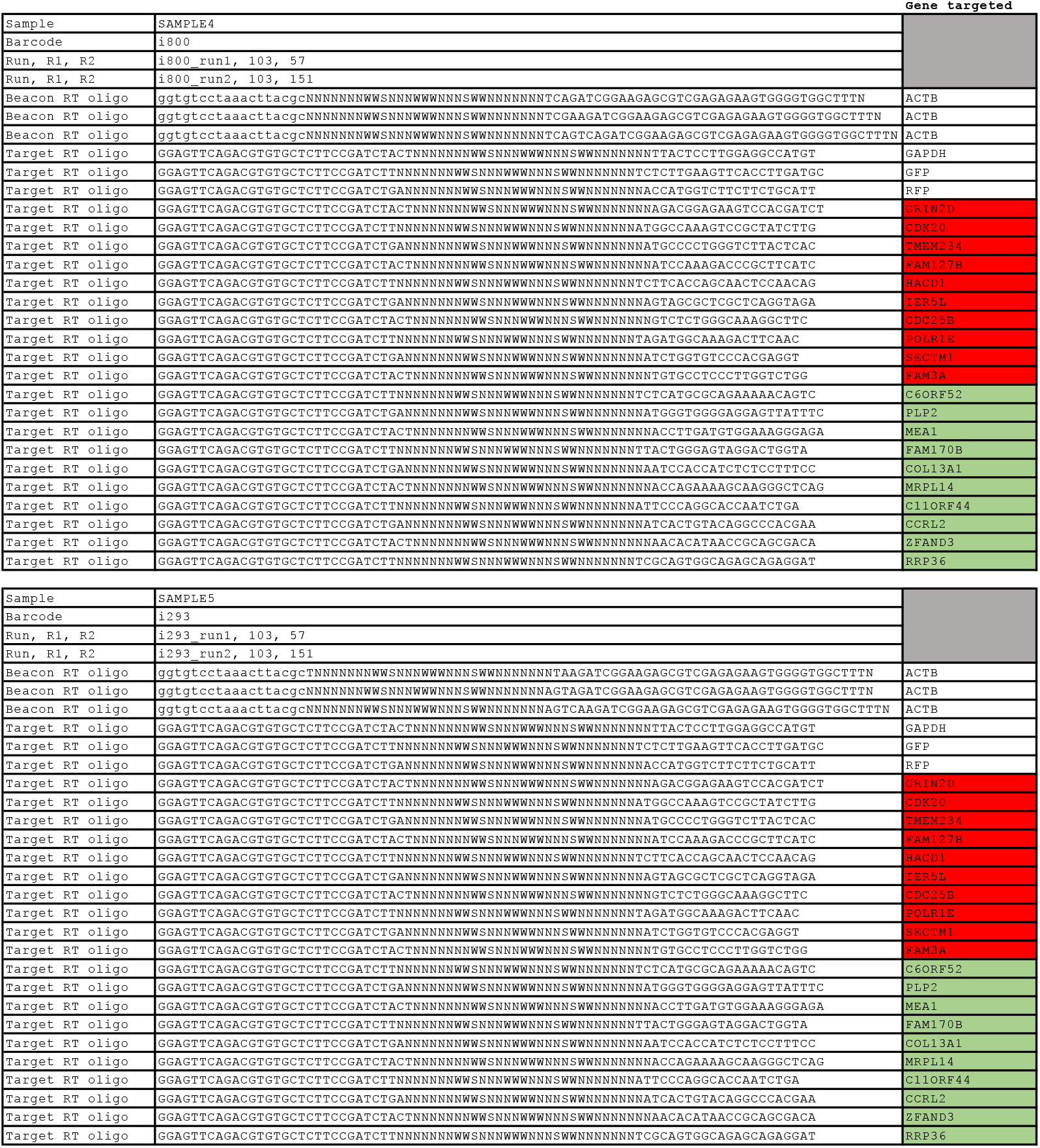
Oligonucleotides used for 24-plex samples during reverse transcription, related to Fig. S5. Genes previously found (http://amp.pharm.mssm.edu/Harmonizome/, CCLE Cell Line Gene Expression Profiles) enriched in BT-549 cells are highlighted red, and genes previously found enriched in MDA-MB-231 cells are highlighted green. Lower case nucleotides indicate sequence areas during read parsing for which a 6% error rate is accepted, whereas upper case nucleotides afford zero error tolerance. Read-lengths labeled R1 (beginning at the 3’ end of the beacon UMI) and R2 (beginning at the 5’ end of the target UMI) are shown. All reverse transcription oligonucleotides were obtained as ultramers from IDT Inc.

**Table S5:**
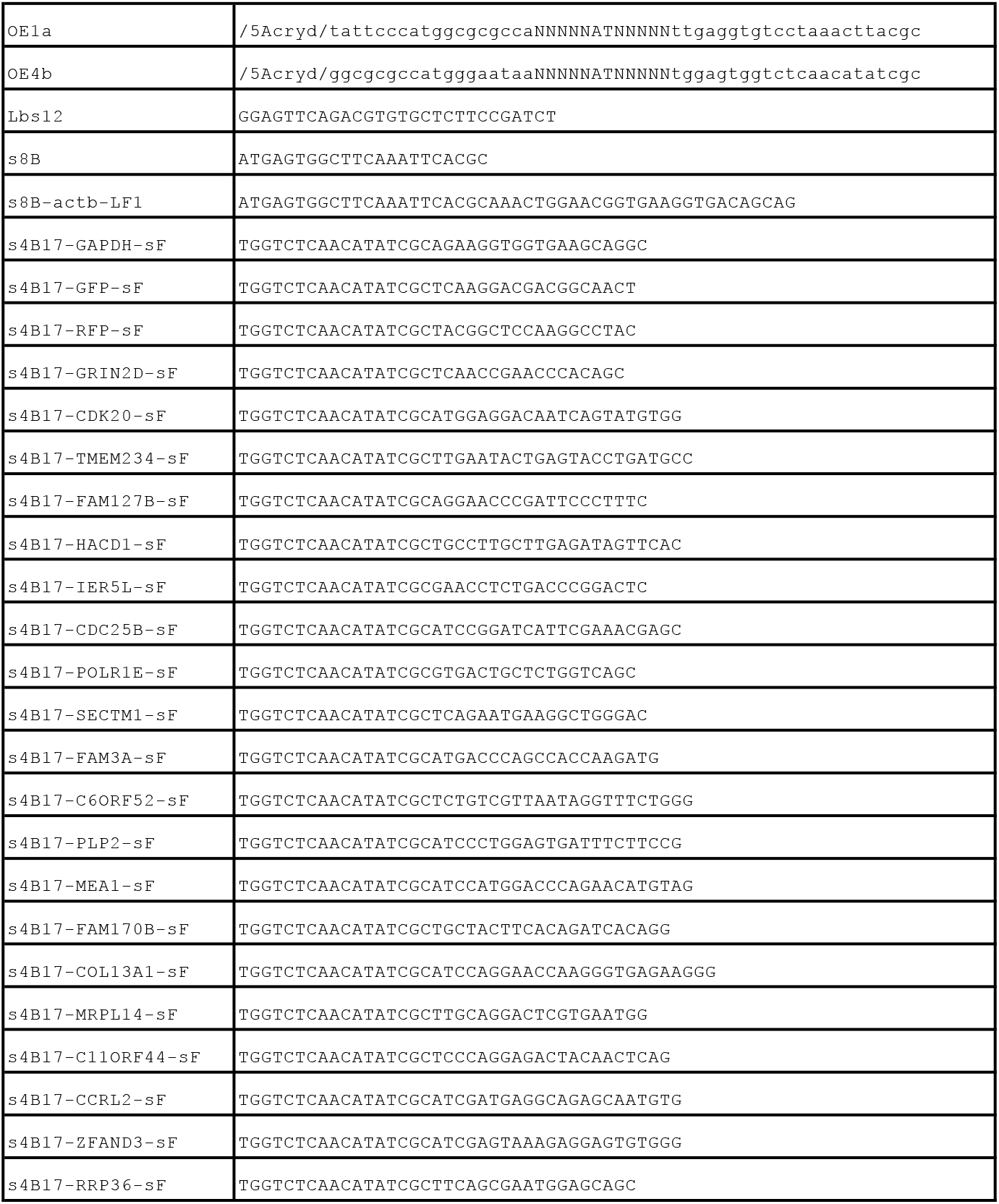
Oligonucleotides used for 24-plex PCR, related to Table S4 and Fig. S5. For PCR, as indicated in the Experimental methods section, gene-specific primers (each ending here in “sF”) were used at a final concentration of 10 nM each (in contrast to 30 nM each for the oligos in Table S2 used for 4-plex PCR). 5’-acrydite modified oligonucleotides were HPLC-purified by the manufacturer.

